# Dynamics of hydraulic and contractile wave-mediated fluid transport during *Drosophila* oogenesis

**DOI:** 10.1101/2020.06.16.155606

**Authors:** Jasmin Imran Alsous, Nicolas Romeo, Jonathan A. Jackson, Frank Mason, Jörn Dunkel, Adam C. Martin

## Abstract

From insects to mice, oocytes develop within cysts alongside nurse-like sister germ cells. Prior to fertilization, the nurse cells’ cytoplasmic contents are transported into the oocyte, which grows as its sister cells regress and die. Although critical for fertility, the biological and physical mechanisms underlying this transport process are poorly understood. Here, we combined live imaging of germline cysts, genetic perturbations, and mathematical modeling to investigate the dynamics and mechanisms that enable directional and complete cytoplasmic transport in *Drosophila melanogaster* egg chambers. We discovered that during ‘nurse cell (NC) dumping’, most cytoplasm is transported into the oocyte independently of changes in myosin-II contractility, with dynamics instead explained by an effective Young-Laplace’s law, suggesting hydraulic transport induced by baseline cell surface tension. A minimal flow network model inspired by the famous two-balloon experiment and genetic analysis of a myosin mutant correctly predicts the directionality of transport time scale, as well as its intercellular pattern. Long thought to trigger transport through ‘squeezing’, changes in actomyosin contractility are required only once cell volume is reduced by ∼75%, in the form of surface contractile waves that drive NC dumping to completion. Our work thus demonstrates how biological and physical mechanisms cooperate to enable a critical developmental process that, until now, was thought to be a mainly biochemically regulated phenomenon.

## Introduction

Fluid flows play an important role in biological development, from the definition of an organism’s body plan (1) and vertebrate organogenesis (2) to cytoplasmic streaming (3) and tissue morphogenesis (4). A widely conserved biofluid mechanical transport process whose dynamics have yet to be uncovered and explained takes place during the development and growth of the egg cell. Across diverse species, oocytes develop within germline cysts alongside nurse-like sister germ cells (5,6). A key juncture in oogenesis occurs when these sister cells transport their cytoplasm to the oocyte prior to fertilization; the oocyte grows as its sister cells regress and die (5–8). Although critical for fertility and early embryonic life, the biological and physical mechanisms underlying this transport process are poorly understood.

*Drosophila melanogaster* oogenesis is a powerful and relevant process for studies of ovarian development. The *Drosophila* oocyte develops within an egg chamber, a multicellular structure comprising a germline cyst of 16 cells that are interconnected through intercellular bridges called ring canals and covered by an epithelium (Fig. 1A) (9–11). Once the oocyte has grown to ∼50% of the egg chamber’s volume, all 15 sister germ cells, called nurse cells (NCs), transport their cytoplasm into the oocyte in a process called ‘nurse cell dumping’; with a diameter of ∼10 μm, the ring canals are large enough to permit passage of most cytoplasmic contents (Fig. 1B; Fig. S1A; Movie S1) (12). It was hypothesized that NC dumping is driven by global cortical contractile forces generated through interactions of non-muscle myosin-2 with actin filaments (henceforth actomyosin). According to this hypothesis, the increased contractility brings about an increase in pressure, causing cytoplasm to be ‘squeezed’ out of the NCs and into the oocyte (13,14). Mutants in the myosin regulatory light chain (RLC), encoded for by the *spaghetti squash* (*sqh*) gene, do not complete NC dumping (15,16), suggesting a critical role for actomyosin dynamics in this process. However, in the absence of time-resolved quantitative data, actomyosin’s role in promoting the complete and directional transport of cytoplasm has remained unclear.

**Fig. 1.**
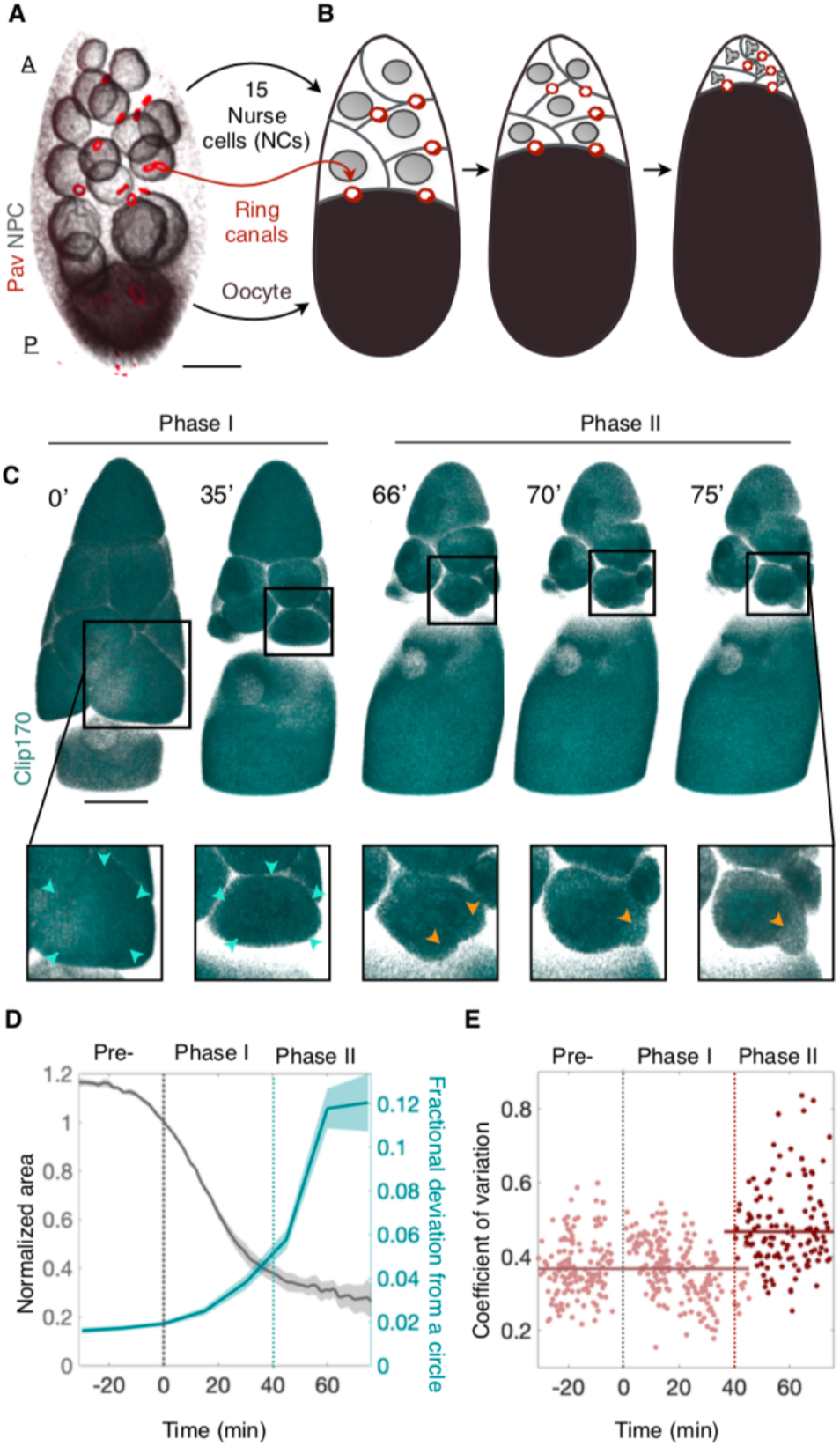
Nurse cell (NC) dumping unfolds in two phases. **A**. 3D-rendered confocal image of an egg chamber comprising 15 anterior (<u>A</u>) NCs (*Nuclear pore complex, NPC*; gray) and one posterior (<u>P</u>) oocyte (black) connected through ring canals (*Pavarotti, Pav*; red). **B**. Schematic illustration of NC dumping: NCs shrink as their cytoplasm flows into the oocyte through ring canals (Fig. S1A). **C**. 3D-rendered time-lapse confocal images of an egg chamber expressing *Clip170::GFP* undergoing NC dumping. Blowups show a nurse cell first shrinking uniformly (cyan arrowheads; Phase I) before undergoing spatially nonuniform shape deformations and bleb-like protrusions (yellow arrowheads; Phase II) that imply increased actomyosin contractility. Scale bar in **A** and **C** = 40 µm. **D**. Quantification of changes in cell size (gray) and changes cell shape (i.e. fractional deviation from a circle; Fig. S2, C and D) prior to NC dumping (Pre-), and during Phases I and II. Onset of non-uniform deformations (dashed cyan line) occurs ∼40 minutes into NC dumping for the samples analyzed (*N* = 4). **E**. Coefficient of variation of cortical Sqh intensity during ‘NC dumping’, showing a transition (dashed red line) from uniform (*N* = 412; Phase I) to non-uniform (*N* = 122; Phase II) distribution at ∼40 minutes into NC dumping, concomitant with onset of dynamic cell shape deformations.

To investigate this process, we used live imaging of egg chambers to reconstruct the collective flow pattern within the 15-cell NC cluster and actomyosin activity during NC dumping. We found that our experimental observations, namely the intercellular directional transport pattern and time scale of this transport phenomenon, in both WT and mutants, are best captured by a networked flow model inspired by the famous two-balloon problem. In contrast to the model of global cortical contraction, our results reveal a different role for actomyosin dynamics during NC dumping: actomyosin surface contraction waves, which have been prominently observed in a variety of biological systems (17–20), allow NC dumping to run to completion. Combined with other recent studies (21), our results demonstrate the importance of hydraulic transport and biological mechanisms in the regulation of multicellular collective behavior during oocyte development in higher organisms.

## Results

Using *ex vivo* live imaging of egg chambers with simultaneously labeled membranes (*Ecad::GFP*), myosin-2 (*sqh::mCH*), and cytoplasm (*Clip170::GFP*), we determined the dynamics of NC dumping and corresponding patterns of actomyosin activity. First, through size measurements of the oocyte and the 15 NCs (Fig. S1A to D), we found that NC dumping unfolds over the course of ∼100 minutes, a period ∼3-fold longer than previously reported through indirect estimates (22,23). We also found that NCs empty ∼75% of their volume into the oocyte through spatially uniform shrinkage of the cells, and in the absence of nonuniform cell shape changes and membrane blebbing that imply contractile force generation (Phase I) (24). In contrast, transport of the remaining cytoplasm (Phase II) is accompanied by dynamic and persistent deformations of NC shape and blebbing (Fig. 1, C and D; Fig. S2; Movie S2).

Similarly, we found that NC dumping onset occurs without changes to cortical myosin level and localization pattern as compared to the previous developmental stage. NCs’ cortical myosin begins reorganizing from a uniform to a nonuniform dynamic cortical pattern (which increases variation of the cortical myosin signal) only ∼40 minutes into NC dumping, coincident with the onset of dynamic NC shape deformations (Fig. 1E; Movie S3; Fig. S3, A and B); no such changes were observed in control membrane markers (Fig. S3C). We conclude that NC dumping unfolds in two distinct phases, only the latter of which coincides with changes in actomyosin distribution and hallmarks of increased actomyosin contractility, such as plasma membrane blebbing (25).

We therefore explored mechanisms whereby directional intercellular fluid flow can occur in the absence of peristalsis-like cell deformations. To that end, we first determined the spatiotemporal pattern of intercellular cytoplasmic transport and found that NC dumping unfolds in a hierarchical manner that mirrors the NCs’ size and arrangement: while the oocyte is the largest cell in the egg chamber, the NCs, which are arranged in four layers (L1-L4; Fig. 2A), also exhibit a descending cell size order according to their distance from the oocyte (26–28). Our data show that NCs directly connected to the oocyte (L1 cells), which are also largest, are first to transport their contents into the oocyte, followed in order by smaller NCs in layers L2-L4 (Fig. 2, B and C).

**Fig. 2.**
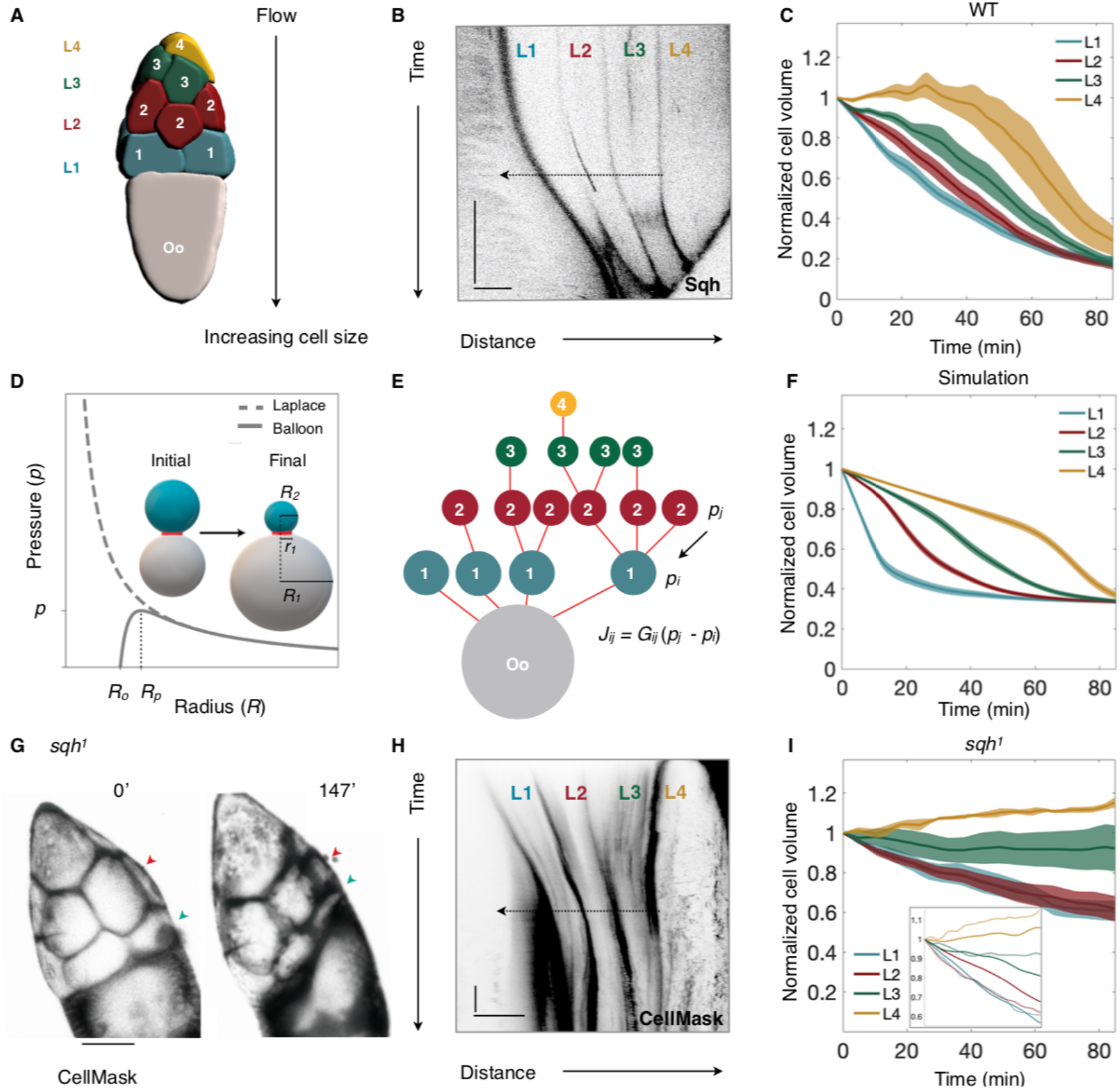
A pressure-driven networked-flow model explains dynamics of NC dumping. **A**. 3D-reconstruction of a germline cyst showing the NCs’ arrangement into four layers relative to the oocyte (Oo). During NC dumping, cytoplasm flows in direction of increasing cell size. **B**. Kymograph of Sqh intensity in WT, illustrating hierarchical onset of NC dumping across the 4 NC layers (L1-L4); arrow indicates direction of flow. Scale bars = 30 min, 50 µm; black indicates highest intensity. **C**. Plot of normalized NC volumes (V/Vo) during NC dumping for each layer from live imaging; *t* = 0 is onset of NC dumping; solid line indicates average; envelopes show standard error (*N* = 15, 12, 9, 5 cells for layers 1, 2, 3, and 4, respectively). **D**. Plots of Young-Laplace’s law and the corrected pressure law for elastic balloons. Pressure is at its maximum, *pmax*, at radius *Rp*; *Ro* is the uninflated balloon radius; *r12* is the radius of the pipe connecting balloons 1 and 2. Schematic illustrates the two-balloon problem, where the smaller balloon (cyan) empties into the larger balloon (gray). **E**. Network representation of the germline cyst in **A** showing cells’ relative sizes and connections; cells are shown as nodes and ring canals as edges. **F**. Plot of normalized NC volumes from simulations of fluid flow in the germline cyst using the best fit parameter set (solid line); envelopes show standard error constructed from the ten nearest sets in parameter space (*N* = 11). Time is scaled by the physical constants of the model. **G**. *sqh^1^* germline mutant showing NCs in the first (blue arrowhead) and second (red arrowhead) layers emptying into the oocyte. **H**. Kymograph of CellMask intensity in *sqh^1^* mutants, showing cytoplasmic transport from the first two layers. Scale bars = 30 min; 70 µm. **I**. Plot as in **C** of normalized NC volumes over time in *sqh^1^* germline clones; (*N* = 14, 17, 7, 6 cells for layers 1, 2, 3, and 4, respectively); inset shows WT cell volume trajectories from **C** (solid lines), re-scaled in time and overlaid with *sqh^1^* mutant data (dashed lines), demonstrating slower yet hierarchical cytoplasmic transport.

Driven by our experimental observations, in which smaller cells empty their contents into larger ones prior to changes in myosin localization and cell shape deformations, we developed a networked fluid flow model that was inspired by the two-balloon problem: if two identical balloons inflated to different volumes are allowed to exchange air, the smaller balloon will empty its contents into the larger balloon that grows at its expense (Fig. 2D). This seemingly counter-intuitive phenomenon can be explained by the Young-Laplace law, which states an inverse relationship between pressure *p* and radius *R* for a sufficiently large balloon; taking into account the hyperelastic behavior of rubber, the pressure inside the inflated balloon is then given by:

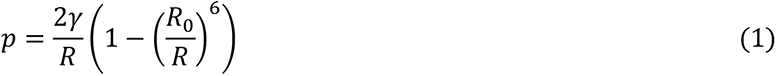

where *R*_1_ is the radius of the uninflated balloon and *γ* is its surface tension (Fig. 2D) (29,30). To apply this description to our biological system, we assumed that the cells are roughly spherical (31) and that cell cortices are incompressible and adequately described by a passive neo-Hookean material under static loading (32) (*Methods*). The two-balloon problem is then readily extended to the germline cyst that can be represented as a 16-cell network (Fig. 2E), where the pressure-driven flux *J_ij_* from cell *j* to *i* through a cylindrical ring canal of radius *r*_ij_ and length *L* is given by:

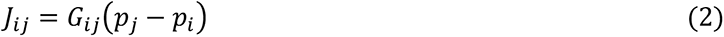

with a hydraulic conductance 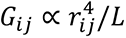 for Poiseuille-type flow (33). The effective cell surface tension in this description arises from the contributions of both in-plane tension of the plasma membrane and actomyosin cortical tension. Using experimentally determined cell sizes at onset of NC dumping as initial conditions (27), we numerically solved the transport equations for the evolution of cell volumes *V_i_* in the 16-cell tree:

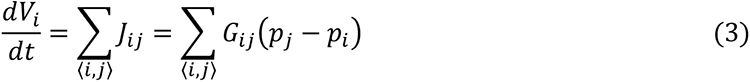

where the sum runs over connected cell neighbors *i* and *j*.

Despite its minimal character, the flow-network model robustly captures qualitatively essential features of the experimentally observed transport dynamics (Fig. S4, A and B; *Methods*). Specifically, the model correctly predicts both the hierarchical pattern of intercellular transport and timescale of NC dumping (Fig. 2F). By also accounting for natural cell-to-cell variability in effective cell surface tension (34), the proposed model successfully captures experimentally observed complex transport patterns along the 16-cell tree. For example, our data show the L4 NC transiently increasing in size during NC dumping, which can occur if the L3 cell to which it is connected shrinks sufficiently such that it becomes smaller than the L4 cell. Such transient back flow away from the oocyte is a feature of NC dumping that has been documented before (35) and is predicted by our model (Fig. S4C, Movie S4).

An insight provided by the model is the high sensitivity of intercellular transport to changes in ring canal size through quartic scaling of the hydraulic conductance. By reconstructing ring canal growth dynamics, we show that ring canals reach approximately 10-fold their initial size throughout oogenesis, consistent with previous studies (12), but also that they do so rapidly prior to the onset of NC dumping (Fig. S5A). It is therefore possible that this an increase in ring canal size sharply accelerates cytoplasmic transport from nurse cells into the oocyte at the onset of NC dumping (Fig. S5B). NC dumping onset is however likely to be affected by other factors as well, such as interactions between the follicle cells and the germline, or external cues.

Our model and the Young-Laplace law predict that lowering of the effective NC surface tension will slow down the rate of intercellular transport but will not affect transport directionality or hierarchical order. Cell surface tension depends on an actomyosin cortex and it can be reduced by inhibiting myosin (36). Therefore, to test how transport is affected by reducing surface tension, we quantified the spatiotemporal pattern of NC dumping in a mutant in the myosin RLC, encoded by *spaghetti squash* (*sqh*). *sqh^1^* mutant germline clones reduce *sqh* mRNA and protein levels by ∼90% (37,38). We found that while Sqh-depleted germline cysts are ‘dumpless’, i.e., do not complete NC dumping (Fig. 2G) (15), the hierarchical transport pattern observed in WT is largely maintained (Fig. 2, H and I; Movie S5). However, NC dumping in *sqh^1^* mutants proceeded more slowly (Fig. 2I, inset). These results showed that myosin contributes to the baseline level of cortical tension, but not to the direction or order of intercellular cytoplasmic transport, which is consistent with myosin activity not changing in Phase I of transport.

Despite the fact that transport of cytoplasm is initiated in *sqh^1^* mutant egg chambers, NC dumping is not completed in these mutants, suggesting that actomyosin and its regulation are important. Indeed, live imaging of egg chambers with labeled myosin (*sqh::mCH*) and actin (*F-tractin::TdTomato* and *Utr::GFP*) demonstrated that actomyosin is highly dynamic during Phase II of NC dumping. Myosin exhibits a diversity of spatiotemporally organized cortical waves, such as colliding myosin wave fronts, rotating cortical bands, and myosin rings travelling between the cell’s poles, which lead to local and dynamic NC shape deformations, rather than isotropic contractions of the entire cell (Fig. 3, A to E; Movie S6). We also found that actomyosin waves in the NCs travel at ∼0.3 μm/s, a speed comparable to that of Rho-actin contraction waves observed in frog and starfish oocytes and embryos (17). Notably, the intercellular pattern of actomyosin wave onset mirrors that of cytoplasmic transport, starting in NCs closer to the oocyte, i.e., those that shrink first, before appearing in NCs further away (Fig. 3F).

**Fig. 3.**
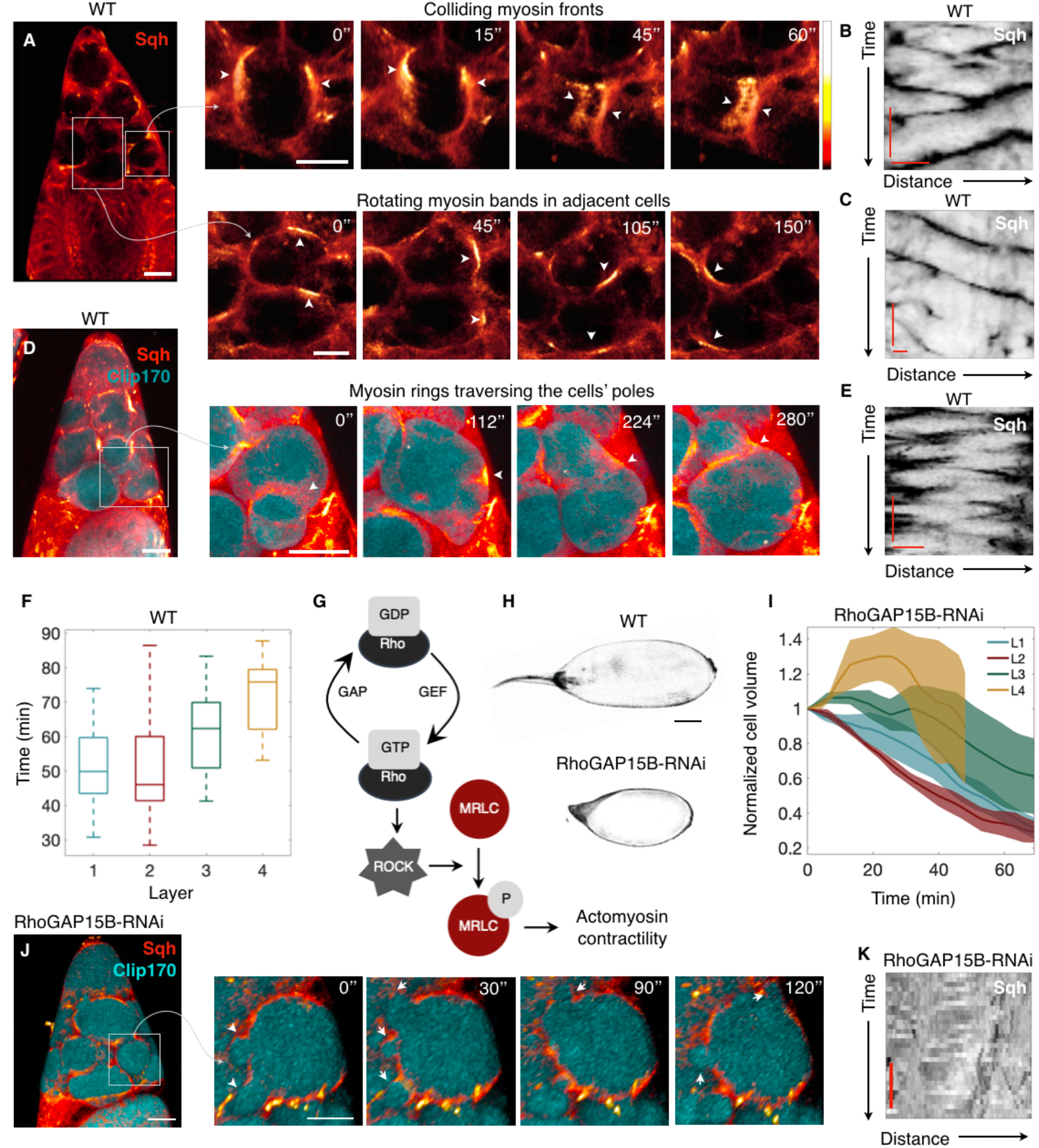
NC dumping completion requires Rho-regulated wave-like actomyosin dynamics. **A**. Heat map of an egg chamber expressing *sqh::mCH*; blowups show NCs with dynamic actomyosin cortical waves as colliding fronts (top) and rotating bands (bottom) in adjacent NCs, with respective kymographs of Sqh intensity around NCs’ perimeter (**B**, **C**). **D**. Heat map of an egg chamber expressing *sqh::mCH* and *Clip170::GFP* (cyan); blowups show a NC with an actomyosin ring (arrowhead) traversing the cell’s opposing poles and deforming cell shape, with kymograph of Sqh intensity in **E**. **F**. Box-and-whisker plot of median time at which nonuniform and persistent cell deformations are first observed following onset of NC dumping in each layer (center line = median; edges = upper and lower quartiles; whiskers extend to extrema; *N* = 20, 22, 16, 5 cells for layers 1, 2, 3, and 4). **G**. The Rho/ROCK signaling pathway regulates phosphorylation of the myosin regulatory light chain (MRLC) and actomyosin contractility. **H**. Comparison between wild-type (WT; top) and RhoGAP15B-depleted (bottom) egg chambers. Scale bar = 50 µm. **I**. Plot of normalized NC volumes during NC dumping for each layer from live imaging of RhoGAP15B knockdowns; *t* = 0 is onset of NC dumping; solid line indicates average; envelopes show standard error (*N* = 7, 6, 3, 2 cells for layers 1, 2, 3, and 4, respectively). The trajectory for the L4 cells stops at *t* ∼ 50 minutes due to membrane breakdown (*Methods*). **J**. RhoGAP15B-depleted germline expressing *sqh::mCH* and *Clip170::GFP;* blowup shows smaller short-lived cell protrusions as opposed to the cell-scale dynamic deformations observed in WT. **K**. Kymograph of Sqh intensity along the perimeter of a cell in a RhoGAP15B knockdown at a comparable time to **B**, **C**, and **E**, illustrating disrupted wave dynamics; black indicates highest intensity. The time scale bar is 5 minutes, while the horizontal axis represents fractional distance along cell perimeter. Scale bar in **A**, **D**, and **J** = 40 µm; scale bar in blowups = 20 µm; kymograph scale bars in **B**, **C**, and **E** = 5 min; 10 µm.

Dynamic actomyosin behaviors like those observed here are regulated through RhoA activation and inhibition (39–41). RhoA is a small GTPase activated by guanine nucleotide exchange factors (GEFs) and inhibited by GTPase-activating proteins (GAPs) (42,43). Binding to downstream effectors, such as the Rho-associated and coiled-coil kinase (ROCK; Rok in *Drosophila*), results in increased contractility through myosin RLC phosphorylation (Fig. 3G) (44–46). Our data show that, similar to myosin, Rok, Utrophin, and F-Tractin exhibit wave-like behavior (Fig. S6, A to C; Movie S7), suggesting that dynamic RhoA regulation, rather than constitutive caspase cleavage-mediated activation of Rok (47), underlies myosin’s dynamics. Furthermore, through a genetic screen (40), we identified RhoGAP15B, the *Drosophila* homolog of ARAP1/3, as a critical regulator of wave dynamics; the RhoGAP domain of the human RhoGAP15B homolog exhibits highest specificity towards RhoA (48), consistent with RhoGAP15B regulating actomyosin contractility. We found that while RhoGAP15B depletion in the germline cyst led to a ‘dumpless’ phenotype (Fig. 3H), onset of NC dumping and the time scale of Phase 1 transport was unaffected (Fig. 3I). The hierarchical pattern of Phase I transport was also largely unaffected, although there was greater variability between the timing of L1 and L2 in knockdown egg chambers (Fig. 3I). Instead, RhoGAP15B depletion disrupted myosin wave dynamics and concomitant cell-scale shape deformations otherwise observed during Phase II in WT: cells displayed erratic myosin activity associated with smaller and more transient cell protrusions (Fig. 3, J and K; Movie S8). We confirmed that incomplete NC dumping in RhoGAP15B knockdowns is not attributable to obstructed or diminished ring canal sizes, or disrupted actin cables (Fig. S7) (49). These data suggest that incomplete cytoplasmic transport in RhoGAP15B knockdowns is due to disrupted wave dynamics in Phase II of NC dumping.

A clue to actomyosin’s role in completing NC dumping came from directly visualizing inter- and intracellular cytoplasmic flow using reflection-mode microscopy. Following actomyosin wave onset and concomitant NC shape deformations, cytoplasm was observed flowing through spaces between NC nuclei and membranes and completing multiple revolutions around the large polyploid nucleus as intercellular anterior-to-posterior transport continued (Fig. 4, A and B; Movie S9). In contrast, intracellular flow in RhoGAP15B knockdowns appeared erratic and lacked the persistent radial motions around NC nuclei necessary for bringing cytoplasm in contact with a ring canal (Fig. 4, C and D). As a result, RhoGAP15B knockdowns exhibited interrupted anterior-to-posterior intercellular flow, repetitive and more frequent transport of cytoplasm away from the oocyte (Fig. 4, E and F; Movie S10), a greater degree of layer 4 expansion (Fig. 3I), and incomplete NC dumping. Given that actomyosin waves first appear in a nurse cell once it has emptied most of its cytoplasmic contents (Fig. S2E), we propose that wave-mediated NC deformations enable continued transport by dynamically creating spaces between plasma membranes and nuclei in shrunken NCs. This creates a transient path allowing cytoplasm to flow past nuclei from anterior to posterior ring canals. Cytoplasmic transport into the oocyte can then run to completion.

**Fig. 4.**
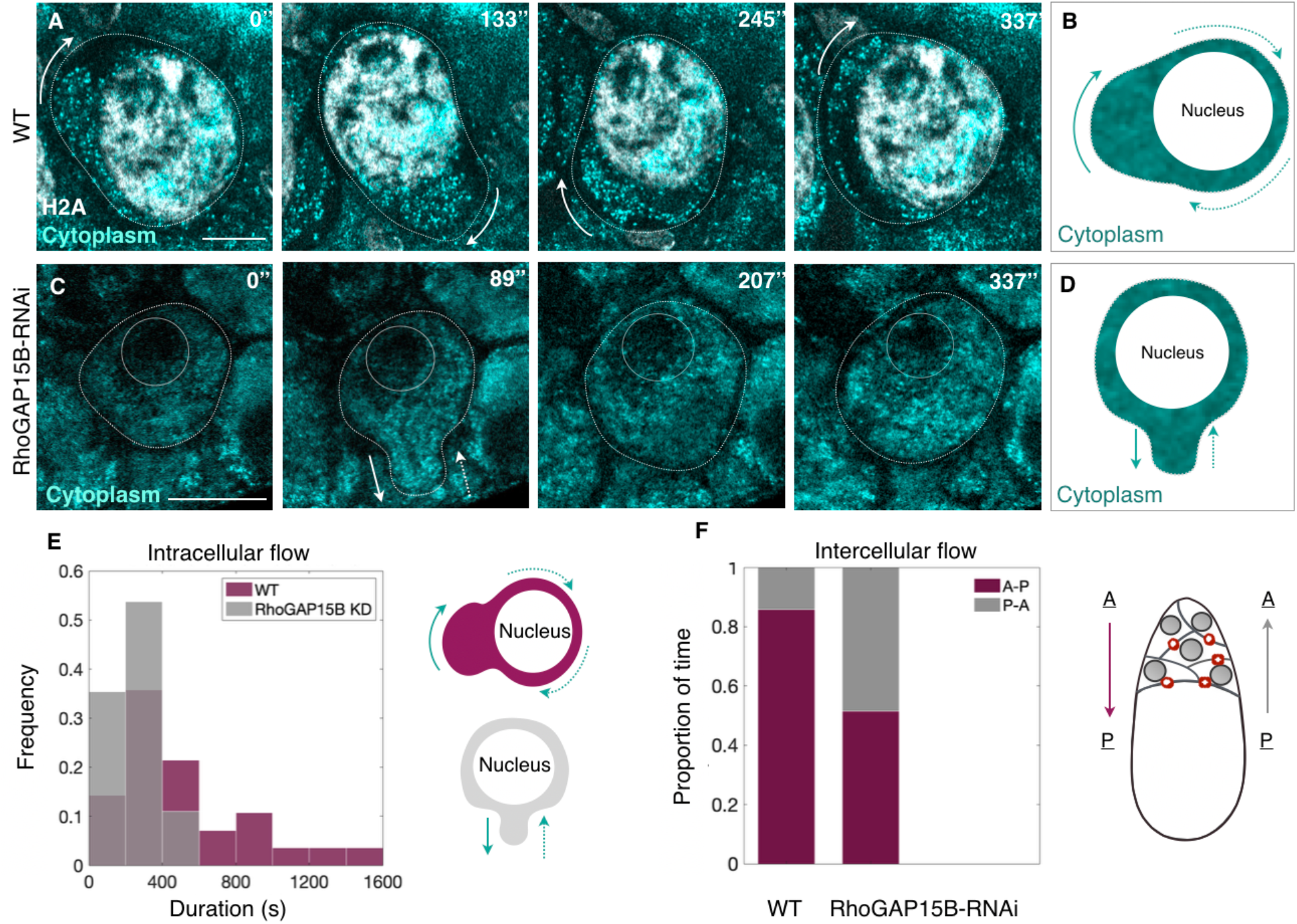
Actomyosin contractions enable continuous intercellular flow in shrunken NCs. **A**. Time-lapse images from reflection-mode microscopy showing cytoplasm (cyan) flowing around a NC nucleus (H2A, white) as persistent actomyosin waves deform cell shape. **B**. Illustration of cytoplasmic flow observed in **A**, where arrows point in the direction of flow. **C**. Erratic and short-lived cytoplasmic flow in a germline RhoGAP15B knockdown, illustrated in **D**, highlighting the lack of repeated revolutionary motions observed in WT. Scale bars in **A** and **C** = 20 µm. **E**. Histogram of the duration of observed intracellular cytoplasmic flow events in WT and RhoGAP15B knockdowns (WT: *N* = 28 events; RhoGAP15B-RNAi: *N* = 82). **F**. Bar plot of the proportion of time anterior-to-posterior (A-P) versus posterior-to-anterior (P-A) flows were observed through ring canals in WT and in RhoGAP15B-RNAi egg chambers (WT: 6 events of intercellular flow spanning 30 minutes total; RhoGAP15B knockdowns: 29 events spanning 54 minutes).

## Discussion

The above analysis presents joint experimental and theoretical work that addresses a longstanding question concerning the origin and regulation of multicellular collective behavior during oocyte development: How do support germ cells in a germline cyst directionally transport the entirety of their cytoplasm into the future egg cell before undergoing cell death? Our experiments and theory show that baseline surface tension and differential cell size provide robust and highly tunable fluid-mechanical control over directional intercellular cytoplasmic flow and that subsequent wave-like actomyosin contractions are essential to complete transport (Fig. S8). These findings contrast with previous hypotheses for NC dumping, which assumed that cytoplasm is driven out of the NCs through a global increase in pressure mediated through upregulated cytoskeletal force generation (29), with increases in actomyosin contractility at the onset of NC dumping (13). Such a model however does not mechanistically explain the directionality and hierarchical flow pattern that is observed during NC dumping; importantly, our data show that changes in actomyosin distribution and concomitant membrane blebbing do not occur until the NCs have emptied most of their cytoplasmic contents into the oocyte. The above analysis thus highlights the complementary importance of physical and biological mechanisms in achieving directed intercellular fluid transport during oogenesis. This study adds to the growing list of examples where hydrodynamics plays a critical role in development (1–4). Lastly, this work also uncovers a diversity of myosin wave-like behaviors and a previously unknown function for excitable actomyosin dynamics as one of the final facilitators of material transfer between sister germ cells, thereby expanding the repertoire of roles played by surface cortical waves in development (17–20).

## Materials and Methods

All experimental data are available in the main text or the Supplementary Information. Experimental protocols are described in the SI Text. The strains and genetic crosses used in this study are in Table S1. We used FIJI and Bitplane’s Imaris to analyze our images and movies, and Matlab for aligning data and generating plots. Measurements and sample sizes are recorded in Table S2. The code used for numerical simulations is available on a public repository at https://github.com/NicoRomeo/balloon-dynamics. A summary of the initial conditions and adjustable parameters is given in Table S3.

### Immunofluorescence and antibodies

Ovaries were extracted through manual dissection. Flies were raised under standard conditions at 25°C and dissected using an established protocol (50). Ovaries from well-fed adult flies were dissected and fixed in 4% (wt/vol) paraformaldehyde at room temperature for 20 minutes and stained with primary antibodies: rat anti-E-cadherin (1:500, Developmental Studies Hybridoma Bank (DSHB)), rabbit anti-PTyr (1:500, Santa Cruz Biotechnology) and mouse anti-NPC (1:500, Abcam). We used goat anti-rat 647nm, goat anti-rabbit 568nm and goat anti-mouse 488nm as secondary antibodies diluted at 1:300. Phalloidin-Alexa-647 (Life Technologies), diluted 1:1000, was used. Samples were mounted in a 50-50 mixture of Aqua-Poly/Mount (Polyscience) and RapiClear 1.47 (an optical clearing medium from SunJin Lab Co.).

### Long-term live imaging

We used a modified version of the Prasad et al. (2007) protocol for live imaging of dissected egg chambers (51). *Ex vivo* culturing of egg chambers took place. Ovaries from ∼2-5 flies were dissected in Schneider’s *Drosophila* media (ThermoFisher SKU #21720-001) that was also used for *ex vivo* culturing; the separated egg chambers were then transferred to a MatTek dish (MatTek #P35G-1.5-10-C) containing 200 μL of media. Imaging took place on an inverted microscope while leaving the lid on the dish to prevent drying. For outlining membranes in the absence of a fluorescent membrane reporter, CellMask deep red plasma membrane stain (Invitrogen #C10046) was added to Schneider’s mix (1:1000).

### Microscopy

Imaging was performed on a Zeiss LSM 710 point scanning confocal microscope with a 25x or 40x/1.2 Apochromat water objective lens, a 10x dry lens, an argon ion, 561 nm diode, 594 nm and 633 HeNe lasers, and the Zen software. Pinhole settings ranged from 1 – 2.5 Airy units. For two-colour live imaging, band-pass filters were set at ∼490 – 565 nm for GFP and ∼590 – 690 nm for mCH. For three-colour imaging, band-pass filters were set at ∼480 – 560 nm for GFP, ∼580 – 635 nm for Alexa Fluor 568, and ∼660 – 750 nm for Alexa Fluor 647. Imaging using confocal reflection mode was performed by setting the detection to the excitation wavelength, i.e. 488 or 561 nm, and using a mirror instead of a dichroic in the beam path. For live imaging of fluorescent reporters, stacks of 20-60 slices and a z-spacing of 2-3 μm were acquired in succession without delay, setting the bottom surface of the egg chamber that faces the cover slip in the MatTek dish or the slide as the last slice, and the first slice as the deepest visible optical plane in the egg chamber. For live imaging using reflection-mode microscopy, a single slice was imaged at a frame rate of ∼0.4-0.9 s/frame in succession. Both modes of imaging took place over the course of ∼0.5-3 hours.

### Genetic screen for RhoGAP dumping mutants

Standard *Drosophila* genetic techniques were utilized to genetically move together tagged proteins or to stimulate transcription of a reporter gene under UAS control through GAL4 expression. This screen was performed to identify maternal effect phenotypes for Rho GAPs, as previously described (40). Briefly, virgins for 22 *Drosophila* RhoGAP shRNA lines were crossed to mat67,15 drivers with fluorescent markers and egg phenotype (i.e., change in egg size) was scored using a dissecting microscope before imaging gastrulation by confocal. UAS hairpin RNA’s were created by the Transgenic RNAi Project at Harvard Medical School (52). Only one of these lines (RhoGAP15B, HMJ02093) gave rise to smaller, ‘dumpless’ eggs.

## Acknowledgements

We thank Stefan Muenster (Max Planck Institute, Dresden) and Wendy Salmon (Whitehead Institute, Keck Microscopy Facility) for advice regarding use of reflection-mode microscopy. We thank Clint Ko and Jaclyn Camuglia for assistance with experiments, and Alexander Mietke for discussions on wave-driven cytoplasmic flows. We also thank Stanislav Y. Shvartsman and Miriam Osterfield for comments on the manuscript. This work was supported by the Molecular Biophysics Training Grant (National Institute of General Medical Sciences, T32 GM008313) (J.J.), a Complex Systems Scholar Award from the James S. McDonnell Foundation (J.D.), the Robert E. Collins Distinguished Scholarship Fund (J.D.), and National Institute of General Medical Sciences grant R01-GM125646 (A.C.M).

## Author contributions

J.I.A. conceptualized the project. J.I.A. and A.C.M. designed the experimental pipeline. J.I.A. and J.J. performed the experiments with input from N.R., J.D., and supervision by A.C.M. J.I.A and J.J. analyzed the experimental data. J.J. developed the computational framework for quantitative analysis of NC dumping dynamics. J.D. and N.R. developed the computational model. N.R. performed simulations, for which J.I.A. and J.J. provided data. F.M. performed the genetic screen that identified RhoGAP15B as an essential GAP for NC dumping. J.I.A. wrote the first draft of the paper. All authors discussed and revised the manuscript.

## Supplementary Information

### This PDF file includes

Materials and methods

Figures S1 to S8

Tables S1 to S3

Legends for Movies S1 to S10

References

### Other supplementary materials for this manuscript include the following

Movies S1 to S10

## Materials and Methods

### Image Analysis

We used FIJI (1) and Bitplane’s Imaris (2) to analyze our images and movies, and Matlab (3) for aligning data and generating plots. Measurements and sample sizes are recorded in Table S2.

### Measurements of cell sizes, volume estimates, and creation of cell volume time courses

Nurse cells are large cells (∼50-70 µm in diameter at onset of NC dumping) and often move noticeably in all three dimensions during NC dumping. As a result, we were unable to consistently capture the entire germline cyst during live imaging, preventing us from obtaining complete cell volume measurements. Instead, we used area measurements as a proxy for volume measurements, assuming cells are roughly spherical and that the volume of a cell could be calculated from its midplane area as 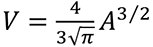. After normalizing to initial sizes, this allowed estimation of normalized volumes as 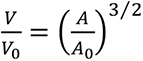, where *A_0_* and *V_0_* are the area and estimated volume at the beginning of dumping for that cell. Areas were measured in FIJI by manually outlining each cell in the *z*-plane of largest cross-section at roughly 5-minute time intervals over the usable course of the movie. We confirmed that area measurements are a good proxy for cell volume by comparing the result of raising our normalized area measurements to the 3/2 power with the few usable normalized volume measurements acquired using Imaris (Fig. S1, B to D).

Nurse cells were not included in the analysis if any of the following occurred before enough data could be taken to generate a trajectory for changes in cell volume during NC dumping: the cell was positioned in *z* such that its cross-sectional area did not show a maximum within the imaged depth, the cell fused with a neighbor early in the acquired time course, or the cell shape was significantly deformed from a sphere when viewed in 3D. Times taken from microscopy metadata were shifted either to align all traces by the time of their dumping onset or to align by wave/deformation onset; that is, to make *t* = 0 correspond to the first frame at which the behavior in question was observed. Because the time required to complete a *z*-stack varied between movies, calculation of mean time courses of normalized volume estimates required temporal alignment. Volume time courses were aligned in Matlab through linear interpolation to 1-minute time intervals. For calculations of layer averages, individual time courses were then truncated to times at which data was collected for all cells to prevent discontinuities in the mean time course resulting from a variable number of measurements. For mean time courses of all cells together, a larger time range was included, and means were produced by grouping data into 20-minute bins and plotting bin averages. Time courses of NC cluster and oocyte lengths (Fig. S1A) were generated using the same methods as for layer averages but using normalized length measurements of the NC cluster and oocyte along the long axis of the egg chamber instead of areas. For the NC cluster and oocyte length time courses, *N* = 4 egg chambers for *t* = −28 to 130 minutes and *N* = 3 egg chambers at other times.

### Dumping onset kymographs and plots

Using maximum intensity projections (MIPs) through *z*, a linear region of interest (ROI) was drawn across the long axis of egg chambers using FIJI, and the MultipleKymograph plugin was used to generate kymographs of myosin (*sqh::GFP* or *sqh::mCH*) or membrane (*Ecad::GFP* or *Gap::mCH*) intensity over time along the line. To generate a plot of average dumping onset rate and progression, several such kymographs were generated, the front between centripetal follicle cells and the NC cluster (i.e. the posterior boundary of the NC cluster) was outlined manually on the kymograph, and its coordinates were taken from FIJI. Coordinates of the fronts were input to Matlab and aligned by subtracting the baseline (initial position of the front). Linear measurements of the posterior NC cluster boundary location showed an S-shaped curve with a clear inflection point, so interpolation was performed as described above and times were shifted to align the inflection in each time course in order to avoid ambiguities in determining dumping onset by eye.

### Time measurements of wave onset

Dumping and wave onset times were collected by viewing movies in FIJI. Time differences between dumping and wave onset were determined, cells were grouped by layer, and layer statistics were calculated using Microsoft Excel.

### Analysis of actomyosin levels

Mean and standard deviation of fluorescence intensity along a segment of the cell perimeter were measured for different time points for germline cysts labeled with either *sqh::mCH* or *sqh::GFP*. Segments were chosen to avoid tricellular junctions, where Sqh intensity was typically much higher than surrounding regions. In addition, the top and bottom 2-3 slices were excluded from the analysis and the outer curves of the nurse cells were avoided to prevent myosin variation in the surrounding follicle cells from affecting the measurements. Note that the ellipsoidal shape of the germline cysts resulted in changes in intensity solely from the amount of tissue between the objective and the optical section; unlike for a flat tissue, this intensity decrease was not consistent across a plane. As a result, segments of the cell perimeter were limited in length and confined to regions without a noticeable intensity gradient along the ROI. Lastly, since a measurement of intensity alone would not differentiate between actual relative biological changes in myosin levels/distribution and imaging artifacts, the coefficient of variation (COV), defined as the ratio of standard deviation to mean (*σ*/*μ*), was calculated for each measurement to determine the fractional variability in intensity along the membrane and allow comparison between different cells and germline cysts (Fig. 1E; Fig. S3B). As a control, the same process was repeated for cells labeled with *Resille::GFP*, a membrane marker (Fig. S3C). Measurements were grouped into three phases, the first corresponding to times prior to wave onset, the second to times at which the membrane showed only uniform shrinkage maintaining cell shape, and the third to times at which the membrane deformed heavily and non-uniformly, changing cell shape. Average values of the COV for each group were calculated by weighting each measured COV by the length of the corresponding ROI. Two-tailed Welch’s *t* tests were performed using Matlab’s ttest2 function with the ‘unequal variance’ option.

### Generation of actomyosin kymographs

For wildtype (WT) wave behavior kymographs, ROIs were drawn on MIP time-lapse images along a curved path following the wave fronts (colliding waves), along the cell perimeter (rotating waves), or across the cell diameter (traveling rings), and wave kymographs showing Sqh levels were generated using FIJI as described above. MIPs through *z* were created using built-in FIJI tools. Cells used for actin behavior kymographs and the RhoGAP15B RNAi myosin kymograph moved significantly over the time period of interest, so ROIs were drawn around the perimeter of MIP time-lapse images, starting at the same location of the cell in each frame. Intensities around these perimeters were measured in FIJI, input to Matlab, aligned and scaled to their fractional position along the cell perimeter, and displayed as kymographs. The RhoGAP15B RNAi kymograph in Fig. 3K is of a layer 4 cell sufficiently long after dumping onset that a WT cell would have begun non-uniform contractions.

### Quantification of cell deformation

Cell outlines were manually drawn in FIJI over MIPs of a cytoplasmic marker (*Clip170::GFP*) or a membrane marker (*Resille::GFP*); their coordinates were taken and input to Matlab, where the cell centroid and projected area were calculated and used to create a circular outline with the same centroid and area as the cell. At each time point, deviations of the measured outline from the circle were then calculated at 180 evenly-spaced angles from the centroid and normalized by the effective cell radius 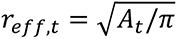 (Fig. S2): for a given angle *θ_n_* at which the projected cell outline is a distance *_n,t_* from the centroid, we defined fractional deformation as 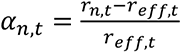 To account for non-circular shapes of cells, which could change over time solely due to volume decrease, the magnitude of change in deformation profile from one time point to the next was then calculated and the root-mean-square (*r_m_s*) value taken:

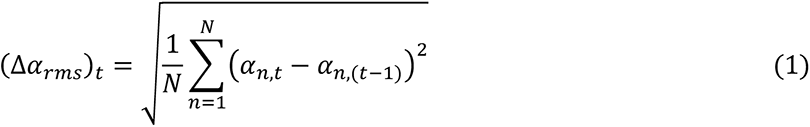

where *n* indexes the angle and *t* the time point; *N* = 180 here. Measurements were aligned in time as described above, so time spacing between elements of *α* was the same for each cell.

### Reflection-mode microscopy flow behaviors

For flow behavior quantification, types of flow were broken into two main categories: intercellular flow (through ring canals) and intracellular flow (induced within a cell by actomyosin waves). For intercellular flow, the direction could be further categorized as either anterior-to-posterior or posterior-to-anterior; the latter is the opposite of that required for complete ‘NC dumping’. For each movie of WT or RhoGAP15B-depleted germline cysts, the duration of each type of flow behavior was recorded. For WT, there were 6 events of intercellular flow spanning 30 minutes total; for RhoGAP15B knockdowns, there were 29 events spanning 54 minutes. There were 28 intracellular flow events in WT for a total of 239 minutes, and 82 intracellular events spanning 338 minutes in RhoGAP15B knockdowns. Fig. 4E represents the actual duration of intracellular events, while the fraction of time undergoing each intercellular behavior was calculated and used to generate Fig. 4F.

### Ring canal growth kinetics measurements

*Z*-stacks of egg chambers dissected at various stages (2 through 11) were acquired as described above. Inner ring canal diameter was measured using Imaris or Zen for every fully visible, undeformed ring canal (26 egg chambers, 10-15 ring canals per chamber, 370 canals total). The stage of oogenesis was estimated using morphological and size features (4) and converted to developmental time (5). Errors in matching measurements of ring canal diameters to a developmental stage within oogenesis arise from inherent variability in developmental rates and uncertainties in determining the exact age of the egg chamber as staging is based on morphological and size features.

### 3. Theory

#### Model

We modeled hydrodynamic flow on the stereotypical 16-cell tree that is the *Drosophila*’s germline cyst. Cells *i* and *j* are connected by ring canals, each of which can be approximated as a small cylindrical pipe of radius *r_i,j_* and length *L*. The viscosity of cytoplasm in *Drosophila* has been estimated *in vivo* (6) as *μ* ≃ 1 *p_a_* ⋅ *s*. At low Reynolds numbers 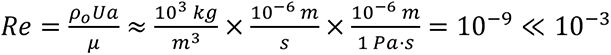, with *U* a characteristic speed, *a* a length scale and *ρ_o_* the density of the cytoplasm), flow through the ring canals is assumed to be incompressible Poiseuille flow. In this case, flow *J_i,j_* from cell *j* to *i* is proportional to the difference in their respective pressures (*p*_j_ − *p*_i_), with the proportionality constant given by the hydraulic conductance *G_ij_*.

The change in the volume of cell is then determined by the total fluxes in and out of the cell:

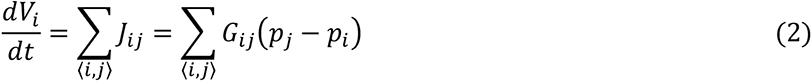

Here, the sum runs over the set of cells connected to cell by ring canals. Summing over all the cells, one can verify that the total cytoplasmic volume is conserved throughout the germline cyst, i.e. ∑*_i_ dV_i_*/*d_t_* = 0 at all times.

#### Balloon pressure law

We model each cell *i* as a spherical ‘balloon’, with interior pressure *p_i_* relative to the homogeneous pressure outside given by the modified Laplace pressure law (7):

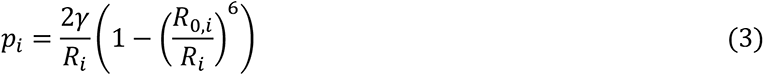

Here, 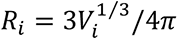 is the radius of the *i*-th balloon, and *γ* is the surface tension of the shell, understood here as the generic energy cost to build an interface. As long as the cell is roughly spheroidal, the spherical approximation will not qualitatively alter the results.

To gain intuition about the behavior of this pressure law, consider two balloons (or two cells) that are connected by an open canal. There are two possible equilibrium configurations for which the pressure in both balloons is equal. In the first case, both balloons are of equal size and the system does not deviate from its initial configuration. In the second case, the balloons assume unequal sizes; that is, an initial size imbalance will be amplified until the pressures balance out. For instance, assuming that initially both balloon radii are larger than *R*_0_, the initially larger balloon will grow to a radius *R_large_* ≫ *R_p_*, whereas the other will shrink down to a radius *R_small_* < *R_p_* where *R_p_* = 7^1⁄6^*R*_0_ ≈ 1.38*R*_0_ is the radius at maximum pressure. Note that as *R*_[_ grows relative to the zero-pressure radius *R*_0,*i*_ one recovers the usual Laplace pressure law, *p_i_* = 2*γ*/*R_i_*.

#### Alternative pressure laws

While the microscopic nature of a nurse cell is much more complex than a rubber balloon, cells can be considered effectively viscoelastic, with a filamentous cytoskeleton bearing similarities to crosslinked semiflexible polymer networks such as rubber (8). Merritt and Weinhaus obtained the above pressure law for the case of an isothermal spherical rubber balloon, using a specific constitutive equation (also called *stress-strain equation*) (7), first derived by James and Guth for rubber in 1943 (9). In fact, as shown previously (10), one would obtain the same functional form for the pressure law for a neo-Hookean material, which itself is a particular case of a Mooney-Rivlin material for small enough extensions. Mooney-Rivlin materials are generally considered to be appropriate models of incompressible rubber-like materials and have been widely used to describe biological tissues, including the cellular cortex (11). If we were to relax the assumption of incompressibility, another family of constitutive relations would follow the Blatz-Ko model (10) and more complex variants. Since cells are usually considered more fluid-like than solid (12), the assumption of incompressibility is reasonable, rendering the Merritt and Weinhaus law an appropriate framework for this problem.

From a more physical viewpoint, to illustrate how a different microscopic model would provide a different power law correction to Young-Laplace’s law, consider for instance a spherical shell of radius *R* consisting of particles that interact according to an repulsive short-range pair potential *V*(*x* − *y*) = −*gδ*^2^(*x* − *y*). Assuming the particles’ surface density is uniform *ρ*(*x*) = *ρ* = *N*/(4*πR*^2^), the energy contribution of this interaction is:

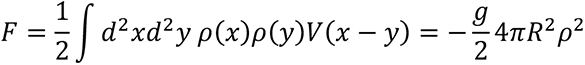

In the presence of a surface tension, the differential of free energy would be, at constant temperature,

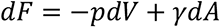

>where *dA* is the variation in area. The pressure as a function of radius would then be given by

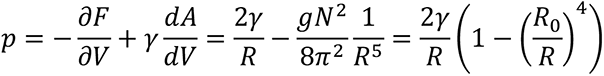

with 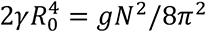. Here we see that the repulsive interaction potential ends up contributing a 1/*R^n^* - a correction to Laplace’s law.

Lastly, to test out whether different ‘elastic’ 1/*R^n^* terms provide qualitative agreement with the data, we ran different simulations with varying values of *n*. While the slopes of curves varied, the directionality and layer-wise hierarchy of dumping order was preserved (Fig. S4B).

#### Hydraulic conductance

Nurse cells in the germline cyst are connected by ring canals. Assuming Poiseuille-type flow (13), the hydraulic conductance *G* of a cylinder of radius *r* and length *L* is given by:

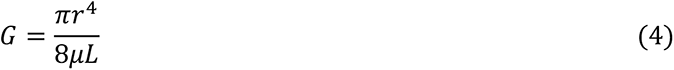

We use this formula to estimate the hydraulic conductance *G_i,j_* between cells *i* and *j* at the onset of NC dumping, adopting typical parameters *r* ≈ 5 *μm* and *L* ≈ 2 *μm* inferred from experimental imaging data.

To test whether such a surface tension-driven flow model could capture the observed dynamics of intercellular networked transport through ring canals, we first estimated the minimal effective surface tension required for Laplace pressure-driven flow at magnitudes comparable to those observed experimentally. The relative contributions of cortical tension versus in-plane membrane tension to cell surface tension are known to vary (14). Here, we combine both contributions into a single effective surface tension parameter *γ* that can be estimated from measurable quantities as described below:

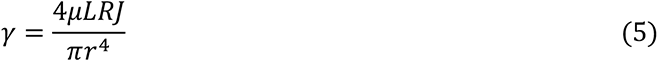

To estimate an upper bound on *J*, the magnitude of flux through a single ring canal, we used the steepest rate of change in cell volume (Fig. 2C), multiplied it by the average volume of a layer 1 cell at NC dumping onset and obtained *J* = 40 *μm*^3^/*s*.

Using *μ* ≃ 1 *pa* ⋅ *s*, *L* = 2 *μm*, *R* ≈ 30 *μm*, *J* = 40 *μm*^3^/*s*, *r* = 5 *μm*, we find a value of approximately 4 ⋅ 10^-3^ mN/m for *γ*. This estimate for *γ* is at the lower end of literature reported values (15), consistent with the known fact that the nurse cell cluster have a relatively low basement membrane stiffness (16). We therefore contend that surface tension is sufficient for driving the observed transport dynamics.

In our subsequent calculations, we assumed that cell surface tension is constant in time for all cells as transport unfolds, which is supported by previous studies showing that plasma membrane tension is actively regulated (e.g. through endocytosis) (17,18). We also assume first that cell surface tension is equal for all cells, justified by the fact that the cortical cytoskeletal network is invariant across cells within the germline cluster and does not appear to be significantly altered during the first phase of NC dumping, i.e., during the phase of surface tension-driven flow.

Additionally, to account for cell-to-cell variability in effective cell surface tension, we explored the effects of fluctuations of surface tension between cells in the model. To this end, we first adjusted the model to the experimental data, and subsequently allowed for approximately 10% surface tension fluctuations to account for variations in shape, basement membrane stiffness and other factors of cell-to-cell variability (see also Section ‘4. Simulations and optimization’ below).

#### Dimensionless equations

Writing out equation (1) fully yields:

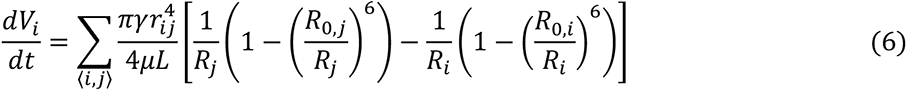

To rewrite this equation in nondimensionalized form we define the characteristic time-scale *τ*_0_ = 4*μL*/(*πγ*) and length scale ℓ_0_ such that the initial oocyte volume is 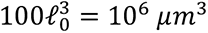, yielding ℓ_0_ ≈ 20 *μm*. Using our estimate for *γ* from above, we find *τ*_0_ ≈ 0.6 s. Defining the dimensionless radius 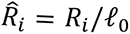, time 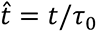, and volume 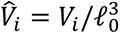, the dynamical equation takes the dimensionless form

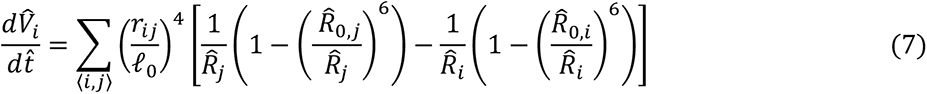

where the dimensionless pressure is given by 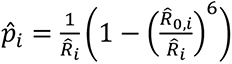.

#### Initial conditions and parameters

At *t* = 0, the sixteen cells have a well-characterized cell size pattern, as nurse cells further from the oocyte are smaller than those that are closer. The initial size distribution is determined as follows (19):

- the oocyte has a volume *V*_0_ equal to the sum of all the nurse cells’ volumes combined
- nurse cells at same distance *d* = 1,2,3,4 from the oocyte have the same cell volume *V_d_* set by
- 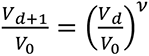 with *v* ≈ 0.84.

Using these initial conditions, we computed the time-evolution of each cell’s volume by numerically integrating the above set of ordinary differential equations. Our simulations (see section 4 below) showed that the trajectories strongly depend on the following parameters:

- Ring canal radii: Assuming that all ring canals at edge distance *d* from the oocyte have the same size *r_d_* (Fig. S5A), we allow for variations of ring canals radii from their corresponding average experimental values by up to plus or minus 60% as described in the section ‘Fitting procedure’ below. Test simulations with equal ring canal sizes do not reproduce the experimentally observed dumping hierarchy.
- Zero-pressure radius: We assume all cells have a zero-pressure radius *R*_0,*i*_ = *ρR_i_*(*t* = 0), where *ρ* is an adjustable parameter between 0 and 1.

Overall, this yields a total of five tunable parameters: the 4 ring canals sizes (*r*_1_, *r*_2_ *r_3_*, *r*_4_) and zero-pressure radius scale parameter *ρ*. A summary of the initial conditions and adjustable parameters is given in Table S3.

### 4. Simulations and Optimization

#### Implementation

The simulations were implemented in Python 3 augmented with the NumPy/SciPy libraries. The regular scipy.integrate.solve_ivp routine was used to solve the system of ordinary differential equations. Simulations were run until the system reached steady state. We have not conducted a full analysis of the stability of the fixed points in a more general setting. However, since our initial conditions are hierarchical, with the oocyte significantly larger than the next largest NCs, the dynamics most likely lie in the basin of attraction of the “1 large; 15 small” solution. Furthermore, those dynamics do not lead to perfectly empty balloons at steady state; indeed, as our experimental data show, completion of the dumping process requires active contractions, which are not included in the network flow model; we can therefore only reliably compare our simulation results with experimental data before the effects of myosin activity become too important, and the simulated steady-state has no physical relevance.

#### Fitting procedure

Experimental measurements provided us with the relative cross-sectional areas of cells, which we averaged per layer of cells at distance *d* from the oocyte. We then estimated the relative experimental volumes 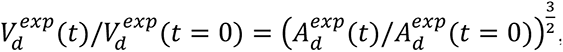, with time *t* expressed in units of *τ*_0_. Our simulations then yield a set of volume trajectories 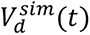 for the average volume of cells in layer *d*.

Since the model developed here describes the first phase of NC dumping, namely, surface tension-driven flow, and does not include the effect of the actomyosin contractility, we compared our trajectories to experimental data taken before onset of actomyosin contractility only. Experiments show that this occurs when cells reach a characteristic fraction of their initial volume *V_wave_* = 0.25*V*(*t* = 0) (Fig. S3). The sum of squares error between the experimental and simulation datapoints was then computed for each of the layer-averaged cross-sectional areas of cells, with a set of weights 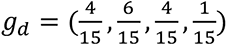 such as ∑_d_ *g_d_* = 1, reflecting the number of nurse cells in each layer. The error is then given by:

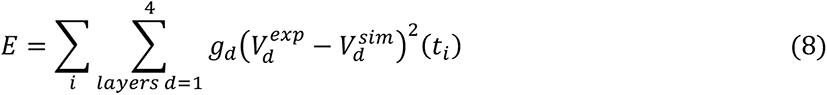

This procedure was repeated as we searched through a grid of possible parameter values around their experimental averages, yielding the error plots shown in Fig. S4A (the final grid search was over 3.2 million simulations). The parameter sets that resulted in the smallest error are denoted as best fit parameters and recorded in Table S3.

#### Cell-to-cell variability

Once the parameters were determined through the method described above, we introduced fluctuations between individual cells in the effective surface tension. To that end, we sampled surface tension for each cell as *γ_i_*= *γ_c_* (1 + *σχ_i_*), where *χ_i_* is a random variable sampled from the standard normal distribution with *σ* = 0.1. Results averaged over *N* = 5,000 trials are presented (Fig. S4C), where the envelope reflects one standard deviation of the fluctuation.

#### Ring canal fits

Measured ring canal diameters (Fig. S5A) were averaged for each layer and each stage. Exponential functions of the form 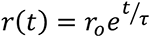 were then fit to the averages using the ‘fit’ function in Matlab. The fit parameters are as follows: *r*_0_ = 0.82, 0.55, 0.44, and 0.67 µm and *τ* = 34.8, 31.2, 29.1, and 38.0 hours for L1-L4, respectively. Values of R^2^ are 0.93, 0.88, 0.85, and 0.83, respectively.

### Figures S1 to S8

**Fig. S1.**
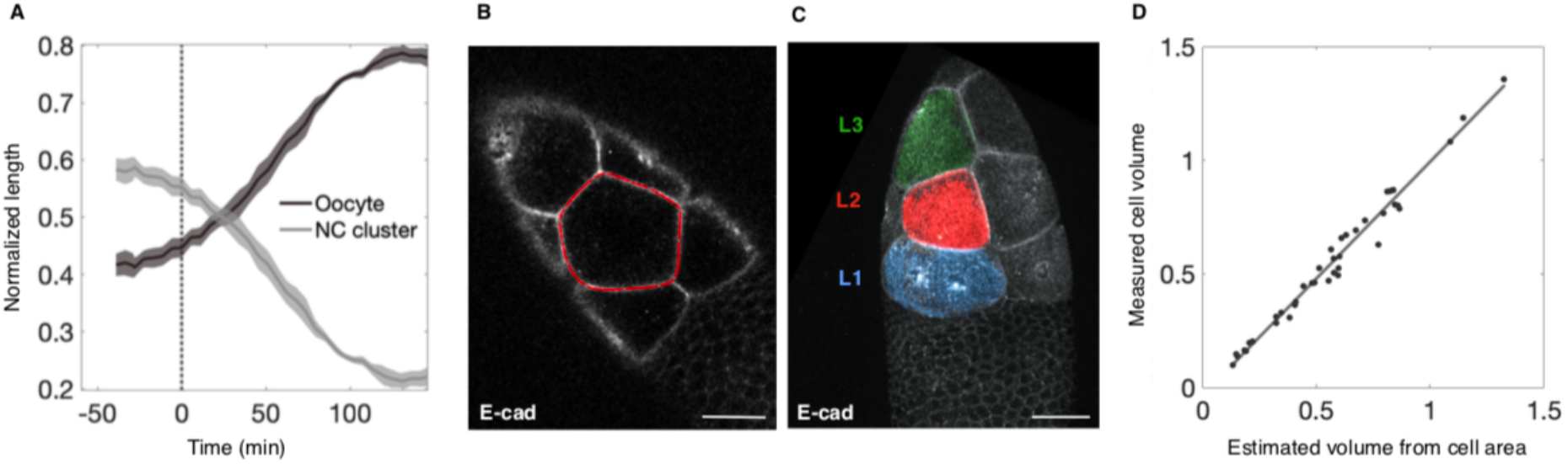
Verification of cell volume measurements. **A**. Plot of fractional lengths of the NC cluster and oocyte during NC dumping (*N* = 4 egg chambers). Line at *t* = 0 indicates onset of NC dumping. **B**. Optical section of an egg chamber expressing *Ecad::GFP*, used to measure NC area as indicated by red outline. **C**. 3D-rendered confocal image of an egg chamber expressing *Ecad::GFP* with surface reconstructions of NCs for measuring NC volume. Scale bar in **B** and **C** = 40 µm. **D**. Plot of normalized NC volume as measured from 3D Imaris surface reconstructions versus normalized NC volume estimated from cell area measurements in FIJI, showing good agreement between both measurements; solid line shows the best fit line; R^2^ = 0.98, with slope = 1.03.

**Fig. S2.**
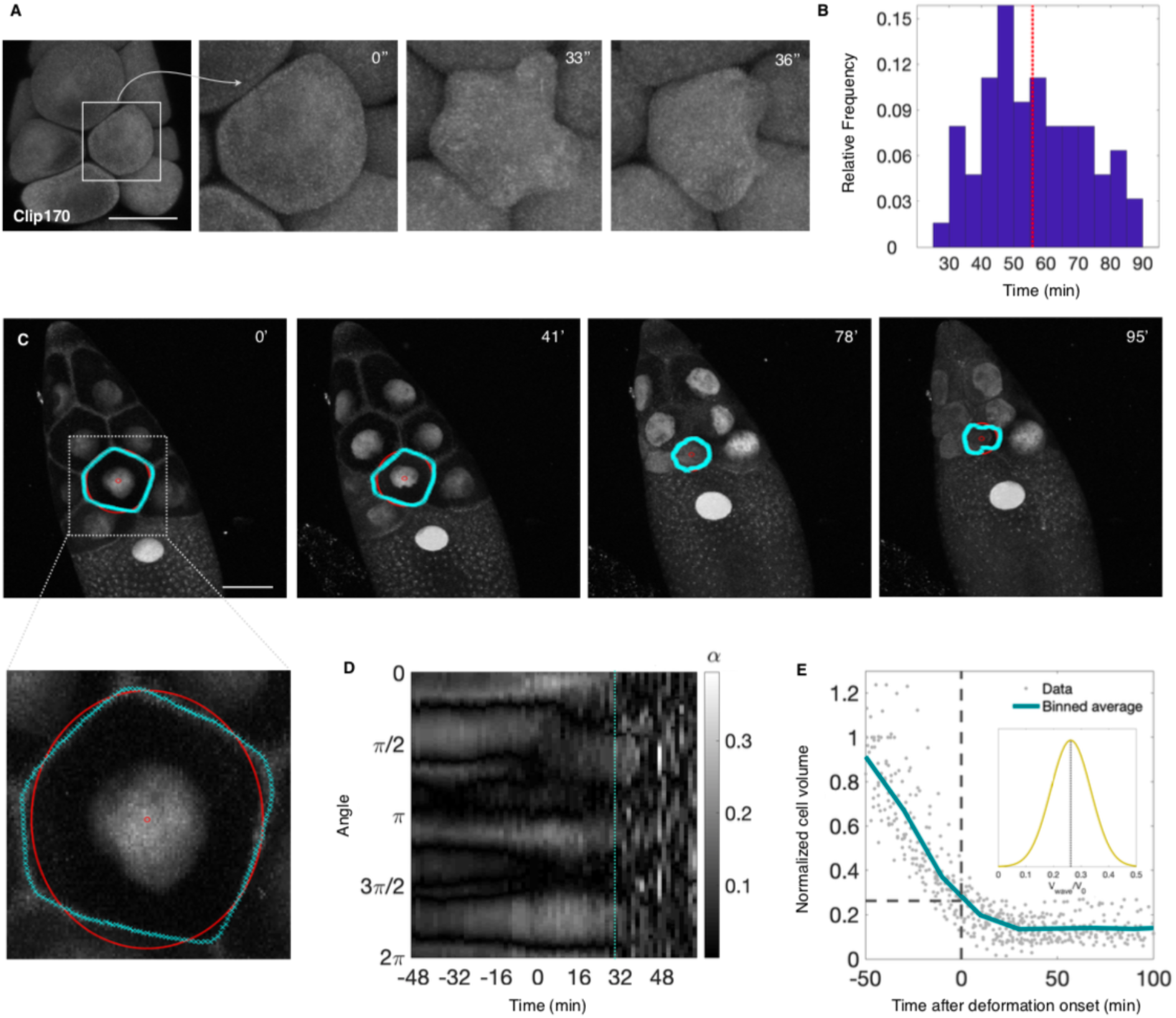
Quantification of nurse cell shape deformations. **A**. 3D-rendered time-lapse confocal images of an egg chamber expressing *Clip170::GFP*; blowups show a cell contracting and deforming nonuniformly with onset of actomyosin waves. **B**. Histogram of time (min) following onset of NC dumping at which nonuniform NC deformations are first observed (red line shows average, *N* = 63 cells). **C**. Time-lapse maximum intensity projections (MIPs) of an egg chamber with labeled membranes (Resille) and nuclei (PCNA); outlined NC is analyzed in **D**. Blowup shows features used to quantify cell shape deformation: red dot indicates the cell’s centroid; red outline indicates a circle with the same projected area as the NC; cyan outline shows point-wise cell shape. Scale bar in **A** and **C** = 40 µm. **D**. Kymograph of fractional deviation of NC shape from a circle, *α*, based on Resille, obtained from the MIP in **C**; line (cyan) marks beginning of nonuniform cell deformations. **E**. NC volume changes of different NCs (*N* = 30 cells) during NC dumping; *t* = 0 defines onset of nonuniform cell deformations. Inset shows the distribution of normalized cell volumes (V/Vo) at onset of cell deformations.

**Fig. S3.**
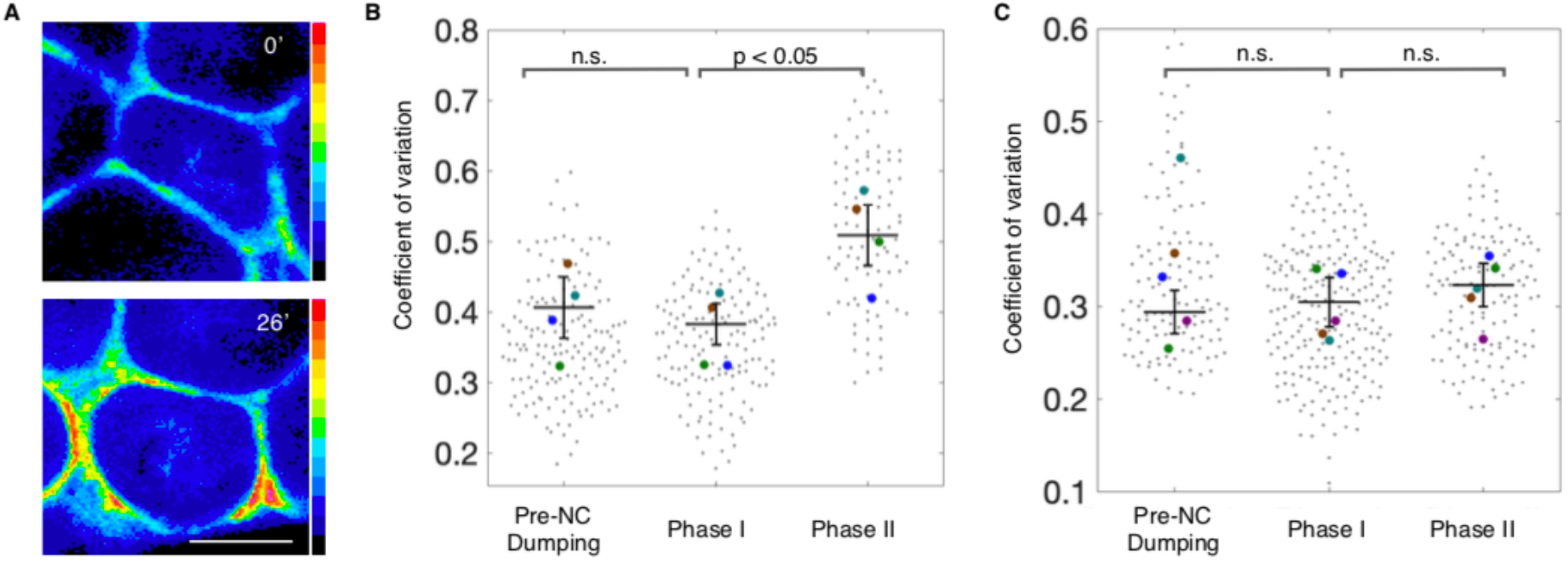
Verification of Sqh distribution measurements. **A**. Heat map of Sqh intensity along the perimeter of an NC in an optical section during NC dumping, showing an increase in intensity and change in distribution. Scale bar = 20 µm. **B**. Swarm plot of the coefficient of variation of Sqh intensity along the cell perimeter in NCs prior to the onset of NC dumping (pre-NC dumping), and during Phases I and II of NC dumping, showing a statistically significant difference in Sqh distribution (*p*-value < 0.05) (*N* = 4 cells; 534 measurements). **C**. Swarm plot as in **B**. showing the coefficient of variation of Resille (a membrane marker) intensity at the cell perimeter in NCs, showing no statistically significant changes in its distribution prior to NC dumping, or between Phases I and II (*p*-value > 0.05; *N* = 5 cells; 504 measurements). Horizontal lines and error bars represent means and standard error, respectively; coloured dots represent mean of each NC.

**Fig. S4.**
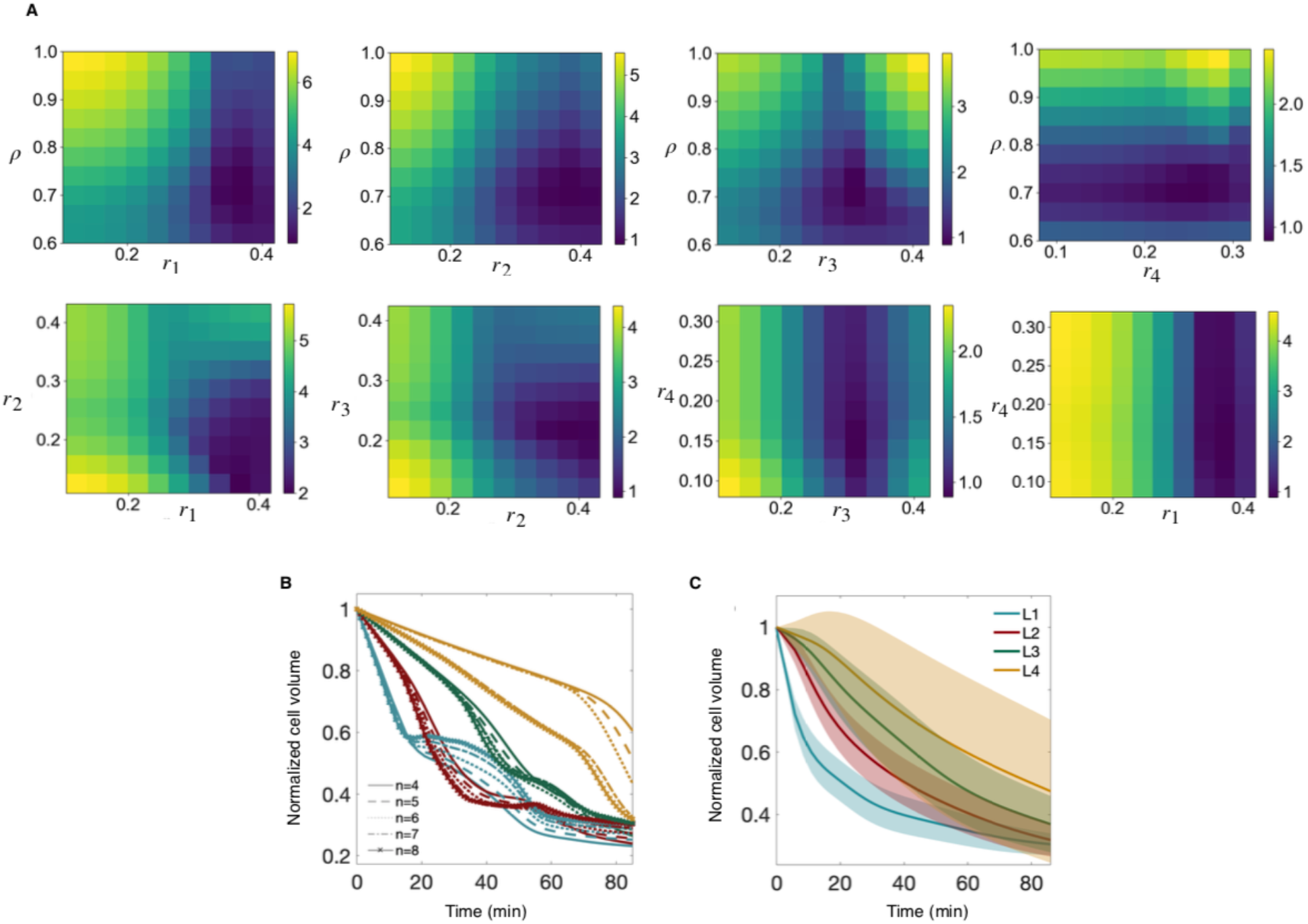
Grid search through parameter space and input parameters of the model. **A**. Two-dimensional slices of the simulation error as defined in *Methods* measured on the 5-dimensional grid space spanned by the sampled parameter ranges (*r*_1_, *r*_2_, *r_3_*, *r*_4_, *ρ*). For a given pair of parameters, the remaining parameters are at their best-fit values. Colour bar refers to simulation error (*σ* = 0.1; see *Methods*). **B**. Results from simulations for varying values of *n*, the exponent in the correction term to the Young-Laplace law, showing that intercellular transport hierarchy and qualitative behaviors are maintained for the values of *n* tested. **C**. Results from simulations averaged over 5,000 trials, where surface tension of each cell is sampled from a normal distribution; envelope reflects half a standard deviation of the fluctuations. As seen in the envelope for L4, in several of these simulations the L4 cell increases in size due to backflow from its downstream L3 cell - a feature of NC dumping that has been reported in previous studies (20).

**Fig. S5.**
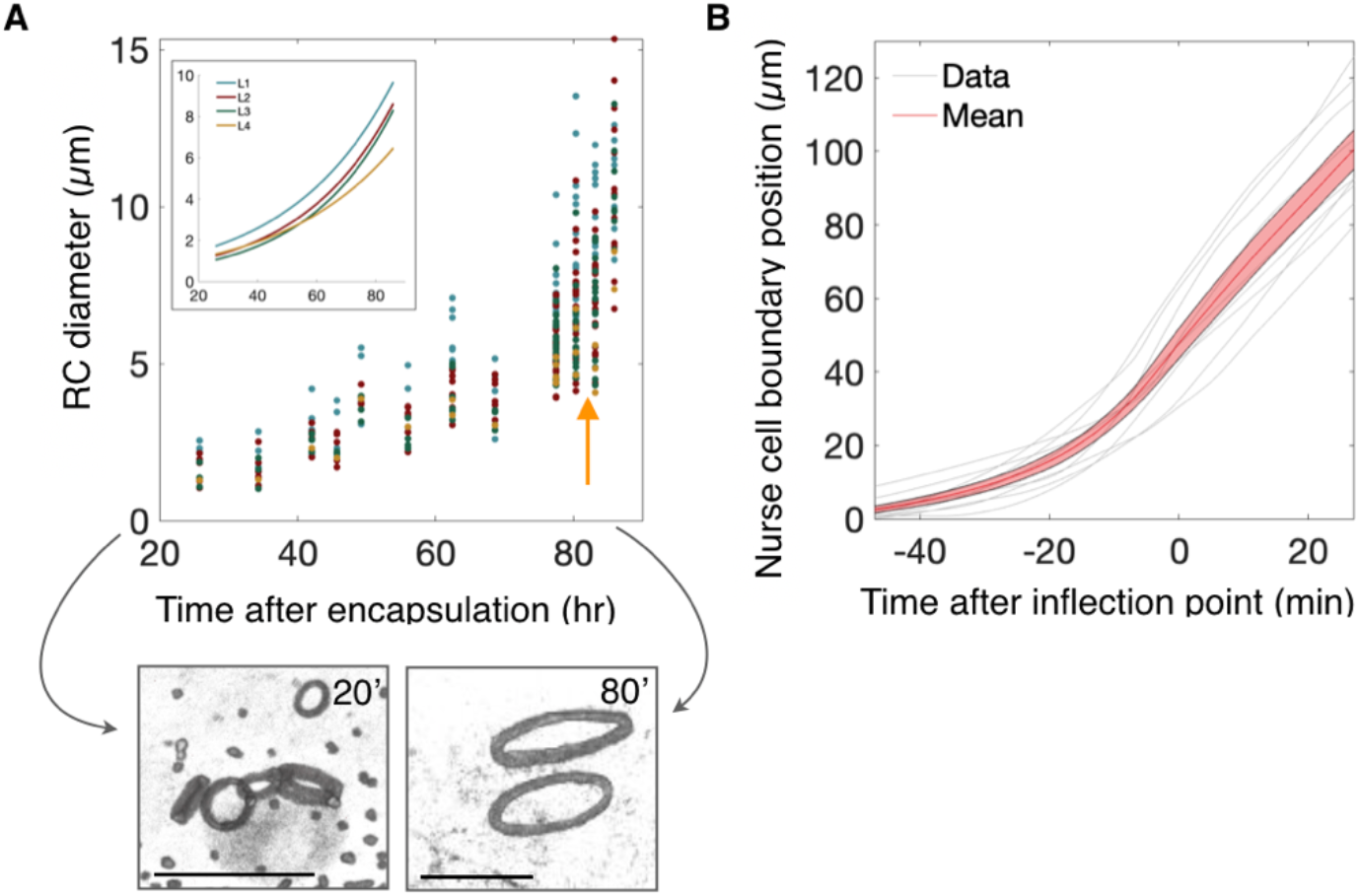
Increase in effective ring canal size correlates with NC dumping onset. **A**. Increase in ring canal diameter during oogenesis, with rapid growth occurring prior to onset of NC dumping (arrow). Inset shows exponential fits to ring canal diameters of the form *d*(*t*) = *d*_0_*e^t/*τ*^* where *d_o_* = 0.78, 0.43, 0.36, 0.64 *μm*, *τ* = 34.8, 31.2, 29.1, 38 hours; *R*^2^ = 0.93, 0.88, 0.85, 0.83 for L1-L4, respectively. Layers are colored as in Figure 2. Arrows point to 3D-rendered confocal images of ring canals at ∼20 and ∼80 hours after encapsulation of the germline cyst (i.e. oogenesis Stage 1). Scale bar = 10 µm. **B**. Measurements of the position of the most-posterior edge of the NC cluster over time, where *t* = 0 is fastest rate of change in position of said edge. Cytoplasmic transport is thus accelerated during NC dumping (envelope shows standard error; *N* = 9 egg chambers).

**Fig. S6.**
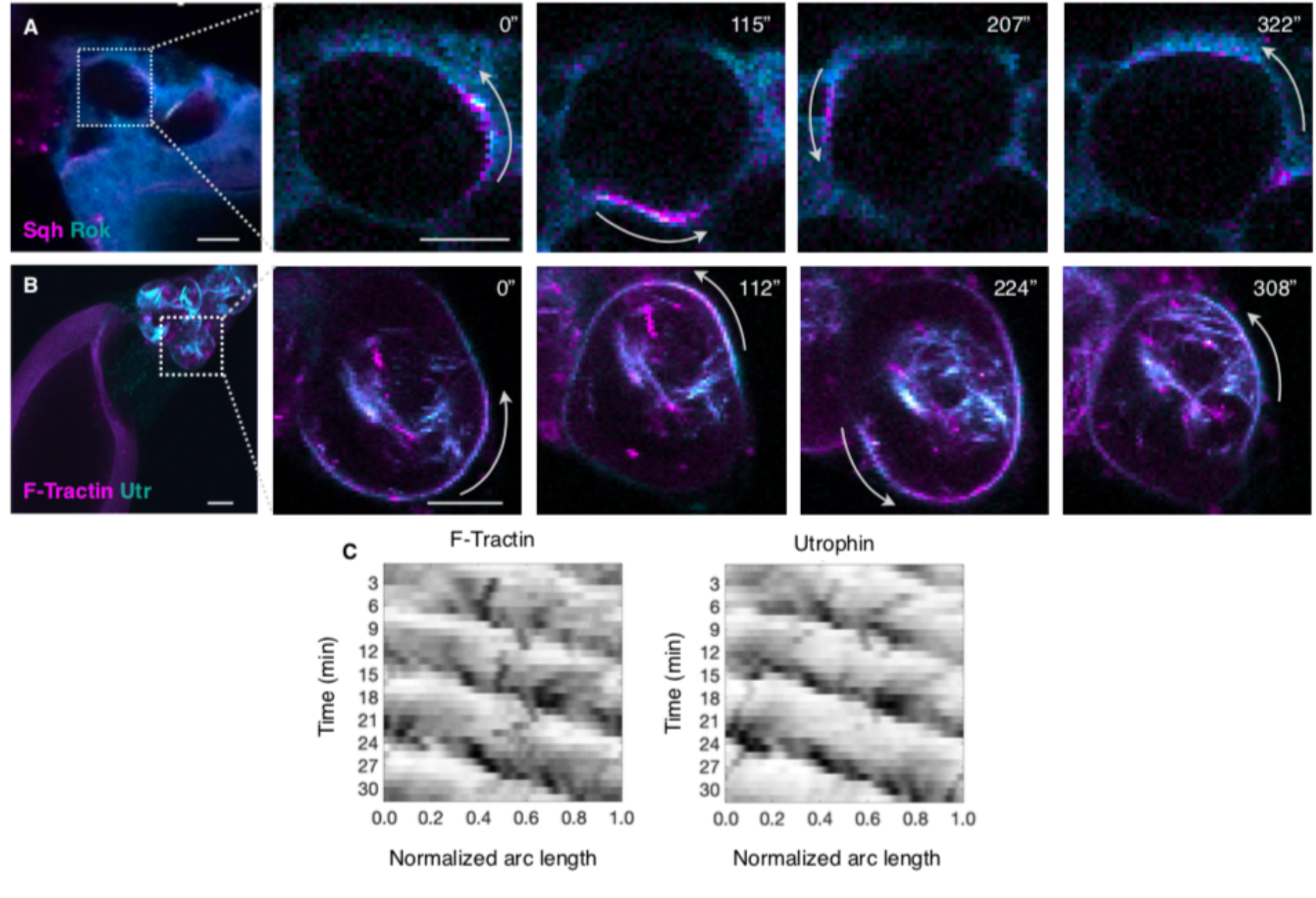
Rho-regulated actomyosin wave dynamics during NC dumping. **A**. Egg chamber expressing s*qh::mCH* (magenta) and *ubi>GFP::rok* (cyan). Rok, upstream of Sqh, exhibits similar wave-like dynamics, seen in blowup as bands rotating along the NC cortex (arrows). **B**. Egg chamber expressing *F-tractin::TdTom* (magenta) and *Utr::GFP* (cyan), with blowup showing both actin-binding proteins exhibiting wave-like dynamics, with bands rotating along the cell cortex (arrow). Scale bars in **A** and **B** = 20 µm. **C**. Kymographs of F-tractin and Utrophin wave dynamics during actomyosin-mediated phase of NC dumping; black indicates highest intensity.

**Fig. S7.**
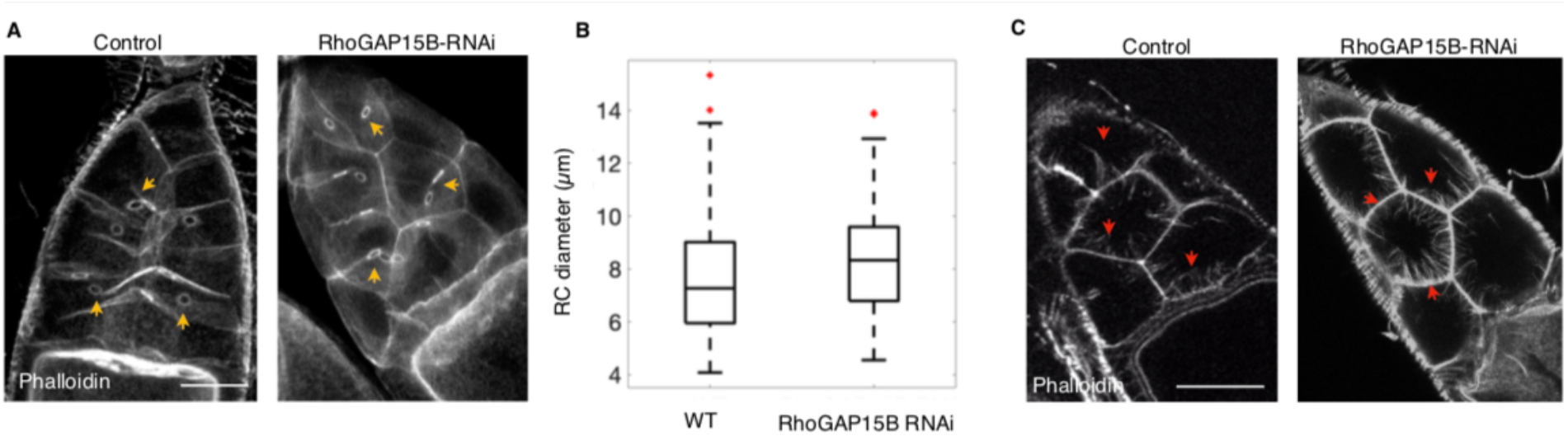
Ring canals and actin cables in WT and RhoGAP15B-depleted egg chambers. **A**. 3D images of a control egg chamber (left) and an egg chamber with a RhoGAP15B-depleted germline cyst (right) stained with Phalloidin to show comparable ring canal structure to WT (arrows). **B**. Box plot of ring canal diameters (µm) in WT and in RhoGAP15B germline knockdowns (*N* = 162 ring canals for WT; 142 for RhoGAP15B-RNAi at oogenesis stage 10 and 11; red pluses mark data points > 1.5xIQR from median). **C**. Optical section of a control egg chamber (left) and an egg chamber with a RhoGAP15B-depleted germline (right) stained with Phalloidin; arrowheads point to actin cables that extend from the cell cortex to tether NC nuclei during NC dumping to prevent their occluding the ring canals. Scale bar in **A** and **C** = 40 µm.

**Fig. S8.**
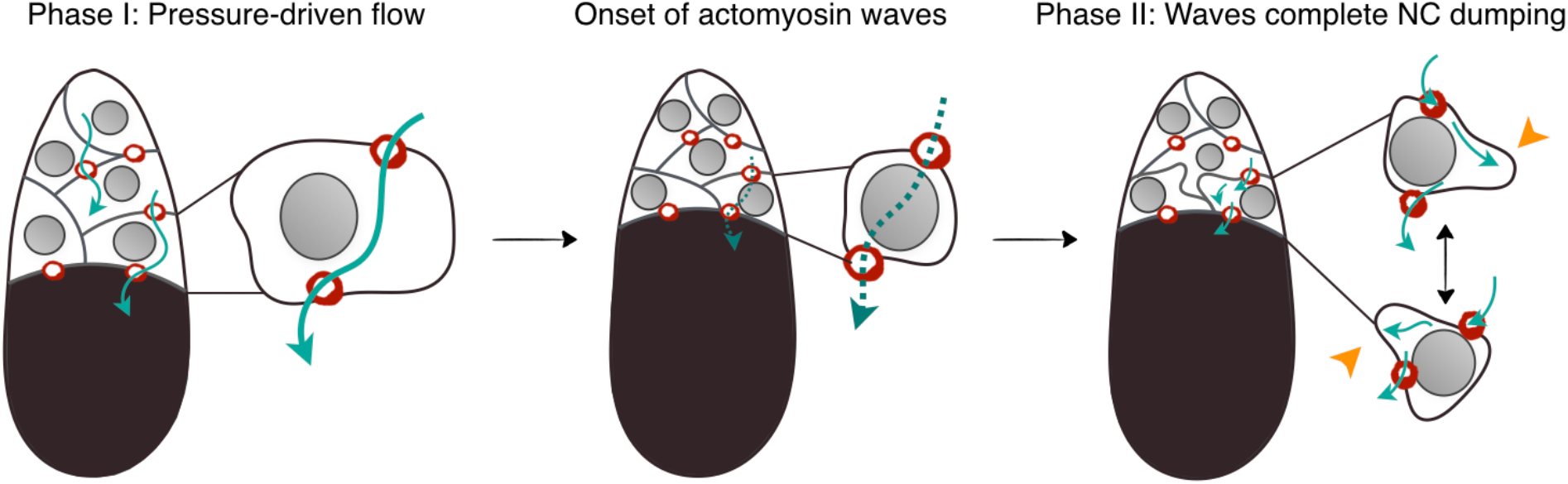
Physical and biochemical mechanisms cooperate to enable NC dumping. Schematic of the proposed model for the contribution of pressure-driven flow with baseline cortical tension and actomyosin-dependent flows to directional and complete NC dumping. Arrows show direction of intercellular flow; dashed arrow indicates interrupted flow; arrowheads point to actomyosin-mediated cell deformations that permit continued intercellular flow in shrunken NCs.

### Tables S1 to S3

**Table S1.**
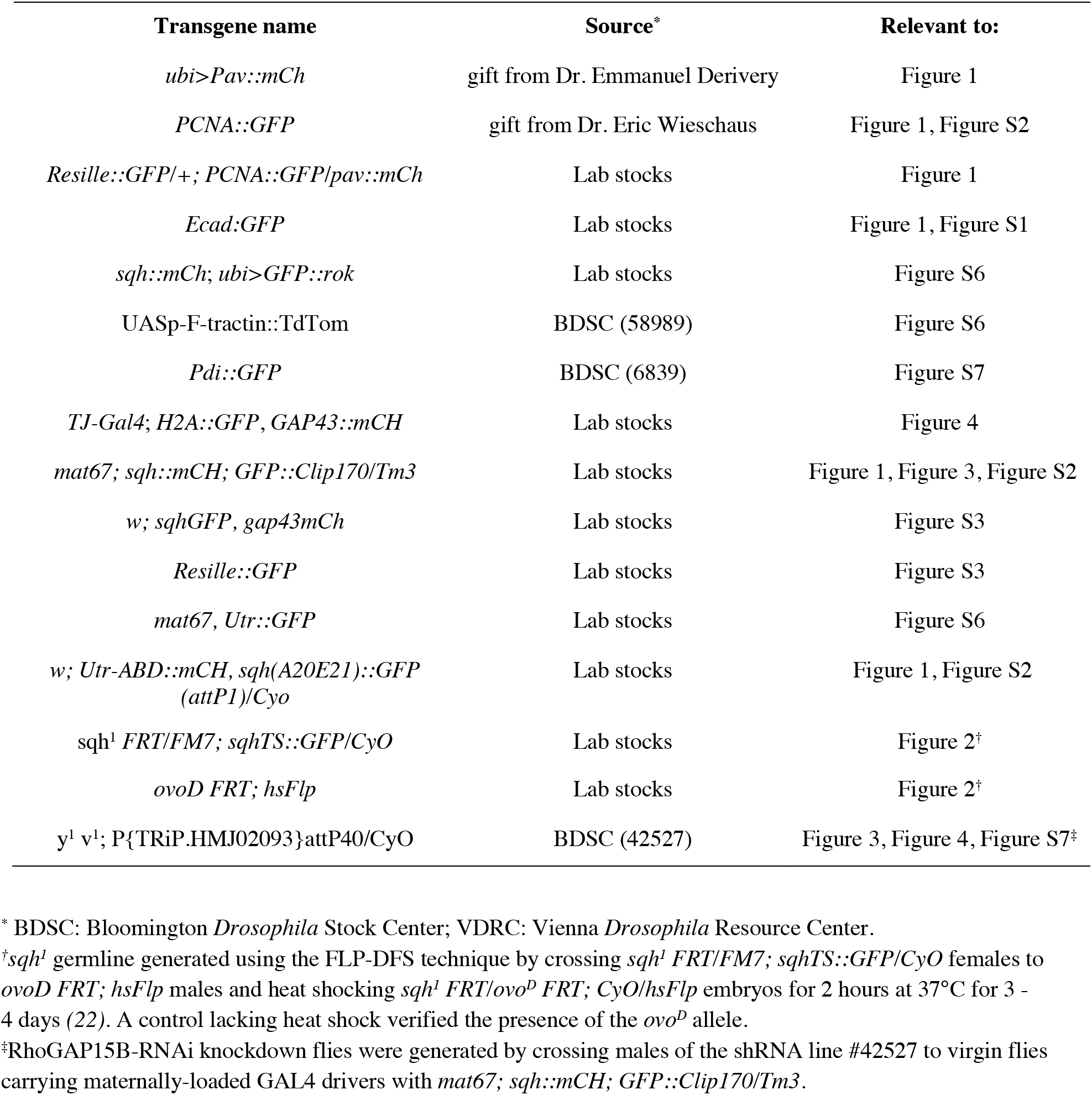
Fly stocks used in this study, their source, and figures in which they appeared.

**Table S2.** Measurement types and sample sizes used to generate the figures.

| Figure | Measurements | N | Number of egg chambers |
| --- | --- | --- | --- |
| 1D | Deformation and projected area | 4 cells | 2 |
| 1E | Myosin COV; points are measurements at different timepoints | 534 measurements from 4 cells | 2 |
| 2C | Cell size over time | 41 cells total | 8 |
| 2I | Cell size over time | 44 cells total | 5 |
| 3F | Wave onset time | 63 cells total | 10 |
| 3I | Cell size over time | 18 cells total | 4 |
| 4E | Intercellular flow behaviors | WT: 6 events spanning 30 minutes;<br>RhoGAP15B: 29 events spanning 54 minutes | 2 (WT), 7 (RhoGAP15B) |
| 4F | Intracellular flow behaviors | WT: 28 events spanning 239 minutes;<br>RhoGAP15B: 82 events spanning 338 minutes | 2 (WT), 7 (RhoGAP15B) |
| S1A | Length of NC cluster and oocyte | 4 | 4 |
| S1D | Cell sizes from FIJI and Imaris | 39 measurements from 9 cells | 4 |
| S2B | Wave onset time | 63 cells total | 10 |
| S2E | Size over time | 30 cells total | 8 |
| S3B | COV of Sqh | 534 measurements from 4 cells | 2 |
| S3C | COV of Resille | 504 measurements from 5 cells | 2 |
| S5A | Ring canal diameter | 370 ring canals | 26 |
| S5B | Nurse cell boundary position | 9 egg chambers | 9 |
| S7B | Ring canal radii at stages 10-11 | WT: 162 ring canals;<br>RhoGAP15B: 144 ring canals | 11 (WT), 10 (RhoGAP15B) |

**Table S3.** Summary of initial conditions and parameters used in the mathematical model.

| Fixed |  |  | Optimized |  |  |
| --- | --- | --- | --- | --- | --- |
| Layer | # of cells | Initial volume* | Parameter | Parameter range<sup>†</sup> | Best fit<sup>‡</sup> |
| Oocyte | 1 | 100 | $\rho$ | [0.22, 1] | 0.55 |
| 1 | 4 | 9.00 | $r_1$ | $[0.11, 0.43]\ell_0$ | 0.37 |
| 2 | 6 | 6.61 | $r_2$ | $[0.11, 0.43]\ell_0$ | 0.40 |
| 3 | 4 | 5.07 | $r_3$ | $[0.13, 0.40]\ell_0$ | 0.32 |
| 4 | 1 | 4.03 | $r_4$ | $[0.10, 0.30]\ell_0$ | 0.26 |

\*In units of $\ell_0^3$ .
<sup>†</sup>Range of ring canal radii is given by the experimental average $\pm 60\%$ for each layer; unitless for $\rho$ .
<sup>‡</sup>Best fit parameters are those that result in smallest error as defined in the methods section.

### Movies S1 to S10

**Movie S1**. Part I: Maximum intensity projection (MIP) of an egg chamber expressing *Ecad::GFP* to mark germline cells’ membranes. The shrinking NC cluster moves anteriorly (top) as cells transport their cytoplasm into the growing posterior oocyte (bottom). Movie length: 4.5 hours; 64-second intervals. Scale bar 50 μm, 1300x real time. Part II: Reflection-mode microscopy of an egg chamber showing intercellular movement of NC cytoplasm (cyan) through ring canals (arrows) towards the oocyte (positioned left). Movie length: 39 seconds, with 0.5-second intervals. Scale bar 20 μm, 10x real time.

**Movie S2**. MIP generated from 4D live imaging of an egg chamber expressing *Sqh::mCH* (red) to label myosin and *PCNA::GFP* (cyan) a to mark the cells’ nuclei. Onset of NC dumping and transport of most of NC volume occurs as NC shrink uniformly without undergoing cell shape deforming contractions. Nonuniform cell contractions commence at 1 hour, 29 minutes (arrow), when ∼75% of NC volume is transported. Movie length: 2 hours 20 minutes; 20-second time points. Scale bar 50 μm, 400x real time.

**Movie S3.** Heat map of an average intensity projection generated from 4D live imaging of an egg chamber expressing *Sqh::GFP*. White line indicates position of NC-oocyte boundary at the onset of NC dumping. As NC unfolds, NC actomyosin cortical pattern changes from a uniform to a non-uniform and dynamic pattern, coincident with onset of contractile NC shape deformations. Movie length: 54 minutes; 20-second time points. Scale bar 50 μm, 480x real time.

**Movie S4.** Animation of NC dumping on the 16-cell network generated using matplotlib in Python, starting with experimentally determined sizes and showing the evolution of cell radii according to the simulation best fit. Ring canals are red when flow proceeds towards the oocyte; blue canals indicate backwards flow. Animation is approximately 2700x real time.

**Movie S5.** MIP generated from 4D live imaging of a *sqh^1^* egg chamber stained with CellMask (magenta). Despite a reduction of ∼90% of Sqh mRNA and protein levels, the first (cyan arrows) and second (red arrows) NC layers nonetheless empty their contents into the oocyte as NC dumping unfolds. Transport is however incomplete, leading to a dumpless phenotype. Movie length: 2.5 hours; 30-second time points. Scale bar 50 μm, 720x real time.

**Movie S6.** MIPs from 4D live imaging of egg chambers expressing *Sqh::mCH* (red) and *Clip170::GFP* (cyan) to mark cells’ myosin and cytoplasm, respectively, highlighting the diversity of actomyosin wave-like behaviors. Left movie shows an example of myosin rings travelling between the cell’s poles (white arrow); right movie shows examples of a colliding myosin wave fronts (yellow arrow) and a rotating cortical band (gray arrow). Movie lengths: 83 minutes with 28-second time points (left); 51 minutes with 17-second time points (right). Scale bar 50 μm; left movie 670x real time, right movie 410x real time.

**Movie S7.** MIP generated from 4D live imaging of an egg chamber expressing *Utr::GFP* (cyan) and *F-tractin::TdTom* (magenta), showing similar wave-like dynamics as myosin. Movie length: 2 hours, 53 minutes; 28-second time points. Scale bar 50 μm, 670x real time.

**Movie S8.** MIP generated from 4D live imaging of egg chamber with a RhoGAP15B depleted germline, expressing *Clip170::GFP* (cyan, left) and *Sqh::mCH* (red, right) to mark cells’ myosin and cytoplasm. RhoGAP15B knockdowns exhibit disrupted actomyosin dynamics and do not complete NC dumping, resulting in a remnant NC cluster and a dumpless phenotype. Movie length: 113 minutes; 35-second time points. Scale bars 50 μm, 840x real time.

**Movie S9.** Left: Live imaging of one slice of an egg chamber, obtained using reflection-mode microscopy to capture cytoplasmic movement (cyan) and stained with CellMask (white) to show cell membranes. As dumping unfolds, cells deform dynamically. Inset: oocyte-NC cluster boundary (yellow line) at NC dumping onset. Movie length: 24 minutes; 10-second time points. Scale bar 50 μm. Right: Blow-up of one slice of a single NC; contractile shape deformations enable continuous cytoplasmic flow past nuclei in shrunken NCs, enabling complete dumping. Movie length: 24 minutes; 10-second time points. Scale bar 20 μm. Both movies are 250x real time.

**Movie S10.** Part I: Live imaging movie showing a single slice of a RhoGAP15B depleted egg chamber during NC dumping, captured using reflection mode microscopy to highlight the NCs’ cytoplasm (cyan). Intracellular flow in RhoGAP15B knockdowns lacks the persistent radial motions around NC nuclei that are observed in WT, instead exhibiting short-lived and erratic protrusions. Part II: reflection-mode microscopy movie of the boxed region highlights rapid reversal of flow direction from normal anterior-to-posterior (A-to-P) to abnormal P-to-A flow through the ring canals (yellow arrow under the ring canal indicates direction). Movie lengths: 170 seconds; 4.6-second intervals, and 298 seconds; 1.4-second intervals. Scale bars 20 μm, 90x real time and 30x real time.

